# Optimizing the hybridization chain reaction-fluorescence in situ hybridization (HCR-FISH) protocol for *Pleurodeles waltl*

**DOI:** 10.64898/2026.04.10.717859

**Authors:** Sofia M. Rebull, Stacy Bendezu-Sayas, Jared A. Tangeman, Erika Grajales-Esquivel, Katia Del Rio-Tsonis

## Abstract

Advances in transcriptomic technologies have transformed the study of complex biological processes, including tissue regeneration, by enabling high-resolution characterization of gene expression programs. In regenerative vertebrate models such as the Iberian ribbed newt (*Pleurodeles waltl*), these approaches can provide critical insight into the molecular mechanisms underlying retina and lens regeneration. However, single-cell and single-nucleus RNA sequencing studies lack spatial resolution, therefore the ability to validate gene expression patterns within ocular tissues is essential and requires optimization.

In this study, we optimized hybridization chain reaction fluorescent in situ hybridization (HCR-FISH) for use in *P. waltl* eyes. HCR-FISH enables sensitive and specific detection of mRNA transcripts through split-initiator probes and hairpin-based signal amplification with automatic background suppression. In addition, because incomplete genome annotation in emerging model organisms complicates transcript selection and probe design, we optimized an optional in silico workflow to support transcript screening, orthology confirmation, and split-initiator probe generation. We systematically optimized fixation duration, proteinase K concentration, and tissue processing parameters to preserve tissue integrity while enhancing signal quality. To overcome imaging constraints imposed by highly pigmented ocular tissues, we implemented a whole-mount protocol with optional bleaching followed by cryosectioning, enabling improved visualization without compromising spatial localization.

Using this workflow, we successfully detected key retinal markers including *SLC1A3* (Müller glia cells) and *RPE65* (retinal pigment epithelium) within the newt eye. Notably, the *RPE65* probe was designed in house and showed comparable detection to a standard Molecular Instruments probe across two sample-preparation protocols. This study presents a reproducible framework for spatial transcript detection in an emerging eye regenerative model and facilitates integration of transcriptomic and anatomical data. Together, the integrated design-to-detection pipeline will strengthen spatial validation of RNA sequencing profiles in *P. waltl*.

## 1. Introduction

Over the past decade, advances in RNA sequencing at single-cell resolution have transformed developmental biology by enabling detailed characterization of gene expression programs that govern cell fate decisions. High throughput techniques such as single-cell RNA-seq (scRNA-seq) and single-nucleus RNA-seq (snRNA-seq) allow for transcriptomic profiling of individual cells to understand what molecular instructions define distinct cell populations (Ding et al. 2020).

In the context of development and regeneration, enormous efforts have been placed toward identifying and understanding the gene expression programs that enable certain organisms to regenerate lost tissues. Newts possess a remarkable ability to regenerate injured organs/tissues such as limbs, heart, brain, lens, and retina (Garza-Garcia et al. 2010); (Kirkham & Joven 2015); (Matsunami et al. 2019); (Tsonis & Del Rio-Tsonis 2004); (Zhang et al. 2002). Of particular importance is deciphering the genetic basis for ocular tissue regeneration because of the significant public health burden that eye diseases pose across the world (National Eye Institute (NEI) 2016). By gaining a deeper understanding of the molecular mechanisms that enable the natural regeneration of eye tissues, it may be possible to generate therapies to combat ocular diseases.

Although unbiased transcriptomic profiling has been implemented in our lab to study lens regeneration in *Pleurodeles waltl*, standard single-cell RNA-seq approaches lack spatial resolution (Choe et al. 2023). Many researchers use immunofluorescent labeling of proteins as a read-out for determining where genes are expressed within tissues. However, immunohistochemistry is often incompatible with non-traditional model organisms such as the newt because these analyses require antibodies that target the protein of interest, and most commercially available antibodies are raised against mammalian epitopes that may not be broadly conserved.

To overcome these limitations, our preferred approach to visualizing gene expression is through direct hybridization of mRNA. Therefore, we implemented hybridization chain reaction fluorescent in situ hybridization (HCR-FISH), a highly sensitive method for detecting mRNA transcripts within intact tissues. With this innovative technology (Choi et al. 2018), RNA probes can be designed for a gene of interest identified in a single cell data set and transcripts can be localized in the native context of the newt eye.

ISH techniques for detecting nucleic acids have existed since the 1960s (Jin & Lloyd, 1997), but the third-generation HCR v3.0 system offered by Molecular Instruments uses split-initiator probes to achieve automatic background suppression (Choi et al. 2018). Each probe carries half of the HCR initiator sequence, which must bind specifically to adjacent regions of the target RNA to form a complete initiator (Figure 1a and 1b). Upon formation of this initiator complex, the first hairpin (H1) is triggered to hybridize, followed by recognition by a second hairpin (H2), resulting in an unbiased chain reaction of alternating fluorophore-labeled H1 and H2 polymerization steps (Choi et al. 2018) (Figure 1a). These self-assembled amplification polymers generate robust fluorescent signal at the site of transcript localization that can then be detected under a fluorescent microscope.

**Figure 1.**
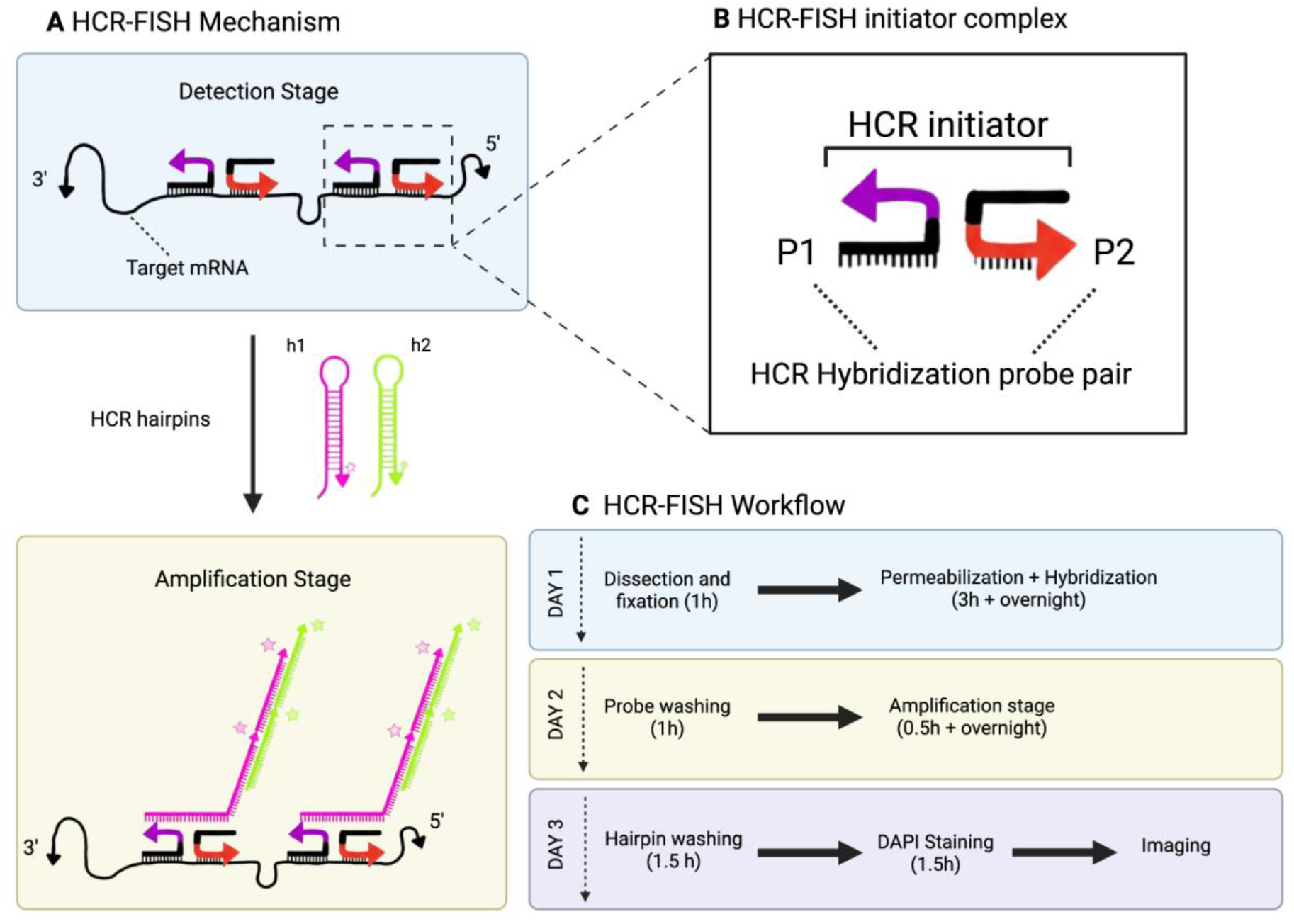
HCR-FISH methodology and overview of workflow. (**a**) During the detection stage, HCR-FISH hybridization probe pair forms an initiator complex and detects the target mRNA. During the amplification stage, the H1 and H2 hairpins (pink and green) recognize the HCR initiator complex and self-assemble into amplification polymers with tethered fluorescent signals (**b**) HCR-FISH hybridization probe pair (purple and red) form an initiator complex. (**c**) General overview of HCR-FISH protocol workflow for *P. waltl* spanning three days. Figure was adapted from Figure 1 in (Huang et al. 2023). *Created in BioRender. Rebull, S. (2026) https://BioRender.com/jrisoom*

Employing this technique for *P. waltl* has presented unique challenges. As an emerging model for studying regenerative paradigms, its genome was recently sequenced, yet annotations remain incomplete. Many novel markers discovered through RNA sequencing lack clear annotation, complicating probe design (Brown et al. 2025); (Elewa et al. 2017). In addition, distinguishing true orthologs from paralogs in a partially annotated genome increases the risk of off-target hybridization when designing split-initiator probes. Furthermore, ocular tissues such as the iris and retinal pigment epithelium (RPE) are highly pigmented, which can interfere with fluorescent signal detection in whole-mount preparations. This has prompted us to explore various approaches for optimizing HCR-FISH in the newt, including depigmentation techniques.

To address the challenges associated with transcript selection and probe specificity, we developed an optional in silico workflow that integrates conserved region screening, orthology confirmation through synteny analysis and paralog discrimination, and an open-source HCR 3.0 Probe design tool to generate order-ready split-initator probe sets. This workflow was validated in *P. waltl* and was experimentally confirmed through successful in situ detection of retinal markers.

Based on protocols provided by Molecular Instruments, HCR-FISH is compatible with whole-mount, formalin-fixed paraffin-embedded (FFPE), and fresh or fixed frozen tissue samples. The whole-mount approach preserves native tissue architecture and avoids sectioning; however, probe penetration and imaging depth can be limited in thick or pigmented samples. In contrast, sectioned FFPE or frozen tissue improves visualization but increases processing time and tissue handling.

In this paper, we present an optimized version for newt eye tissue of the three-day whole-mount protocol developed by Molecular Instruments with a special adaptation for imaging limitations (Figure 1c). This protocol involves processing whole newt eyes with HCR-FISH, followed by a post-hybridization cryosectioning step to circumvent the difficulties of imaging pigmented tissues. As an alternative, we also present an optimized version of the FFPE Molecular Instruments protocol. In both cases we have optimized bleaching approaches.

For researchers seeking to establish HCR-FISH in an emerging model organism, we provide a detailed walkthrough of this protocol in *P. waltl*. By integrating orthology validation, probe design, and optimized tissue processing, this protocol enables spatial validation of gene expression patterns identified through transcriptomic analysis and establishes a framework for mapping molecular dynamics during lens and retina development and regeneration.

## 2. Materials and Supplies

### 2.1 Summary of reagents and equipment

**Table 1.**
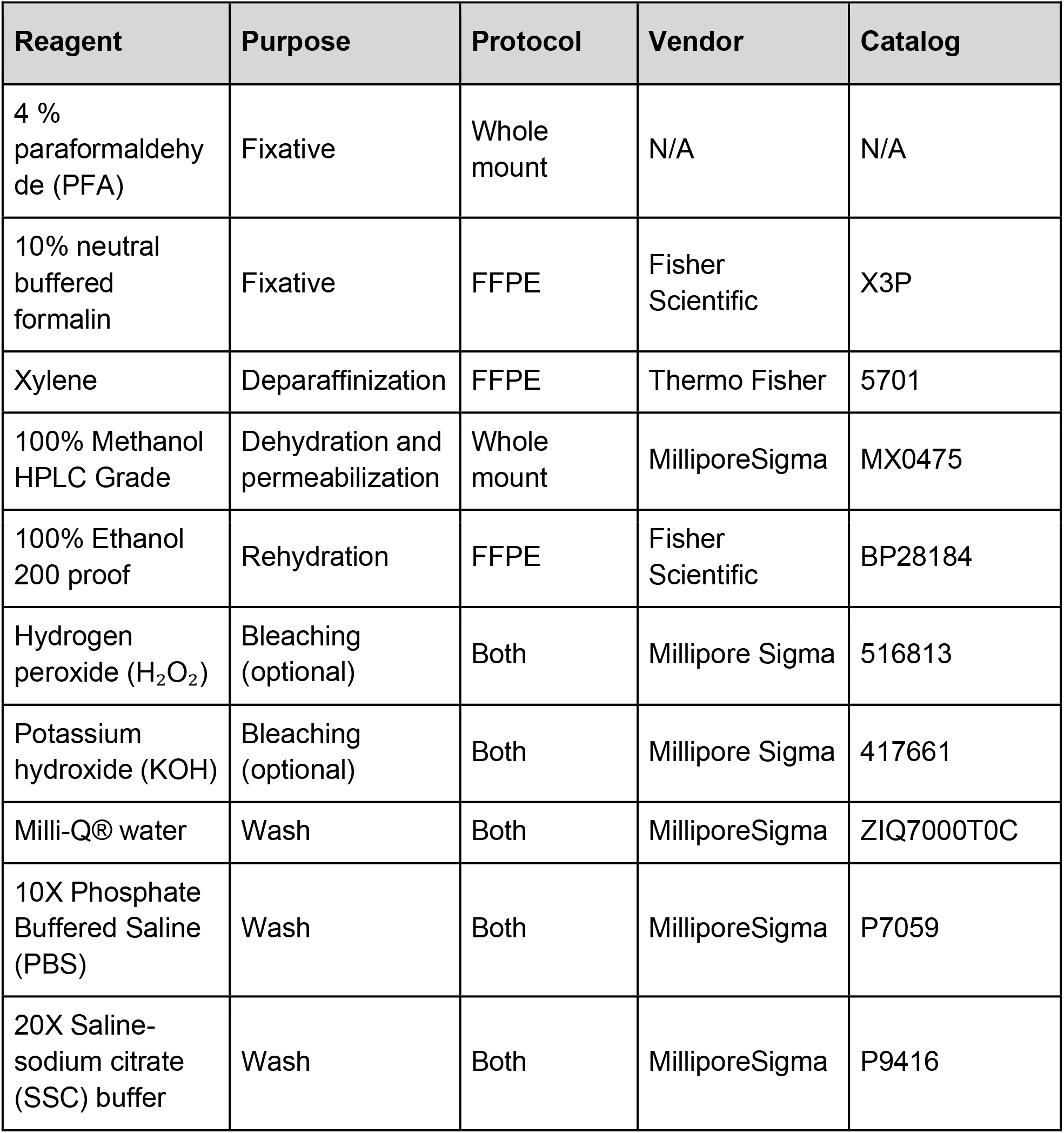
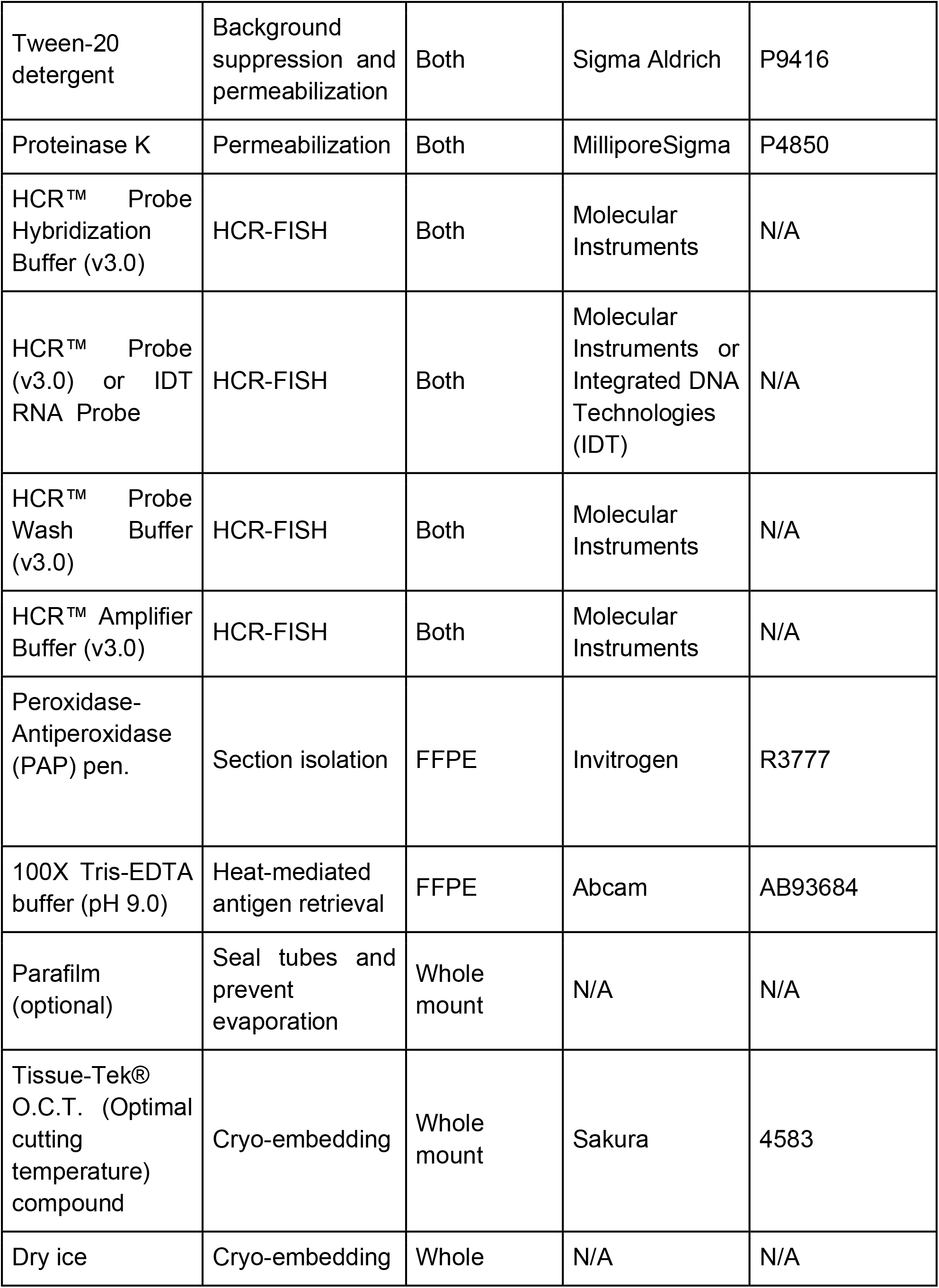
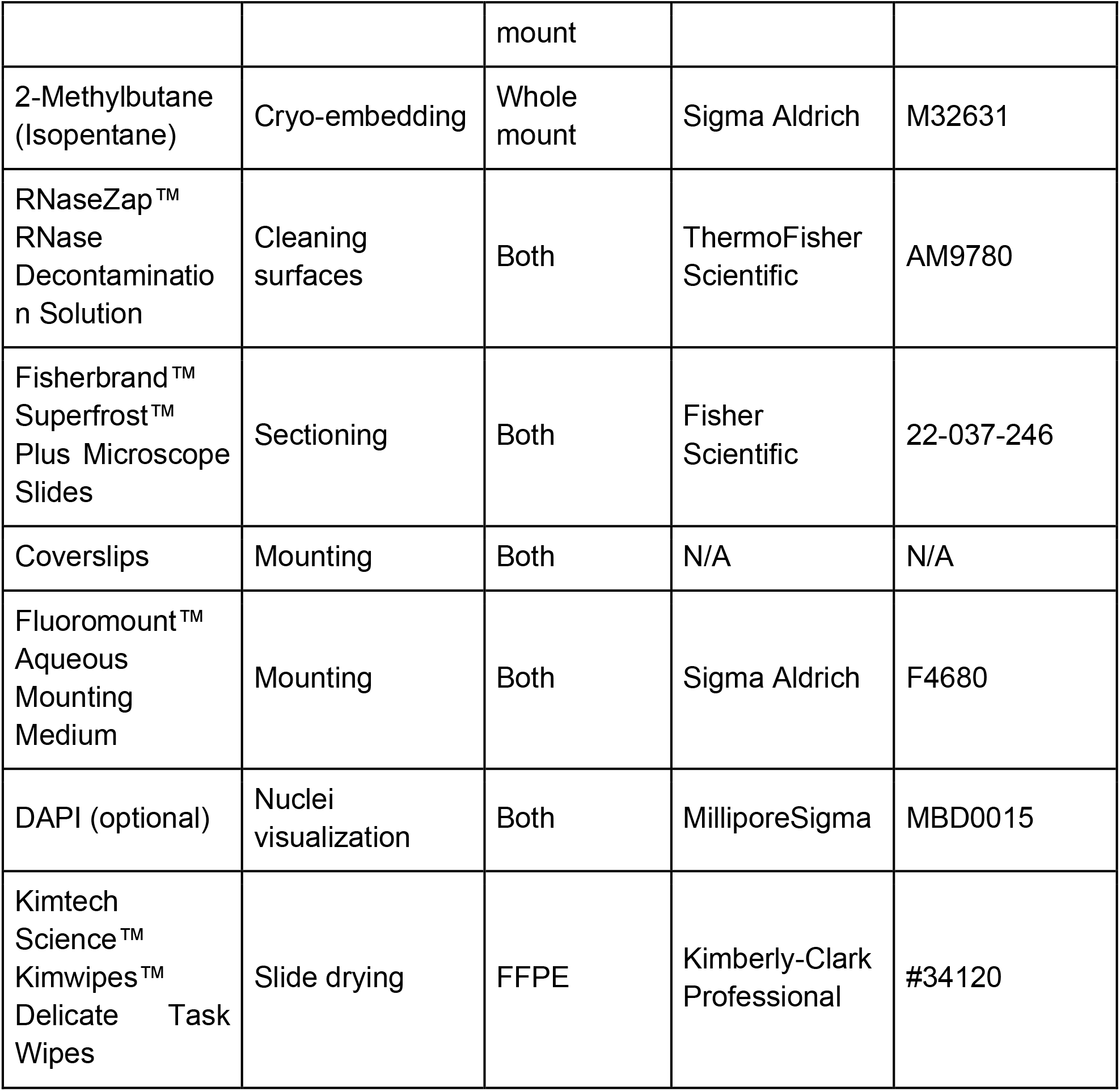
Reagents required for our optimized HCR-FISH protocols in *P. waltl*. All reagents should be molecular biology grade and nuclease-free.

**Table 2.** Equipment used for the optimized HCR-FISH protocol, including vendor and catalog information where applicable.

| Item | Vendor | Catalog |
| --- | --- | --- |
| Fisher Vortex Genie 2™ | Fisher Scientific | 12-812 |
| Dry Bath Incubator | Fisher Scientific | 11-718-2 |
| Multipurpose rotator | Thermo Scientific | 2314 |
| Incubator set to 37°C | N/A | N/A |
| AccuSpin Micro 17 | Fisher Scientific | 75002461 |
| Incubator set to 60°C | N/A | N/A |

### 2.2 Buffer and Solution Recipes

PBS and PBST buffers may be prepared in advance and stored for up to 6 months at room temperature. SSCT buffers may be stored for 3–6 months. All other solutions should be prepared fresh. A ribonuclease-free environment must be maintained throughout the procedure. The use of filtered pipette tips is recommended.

1. 1X PBS for washes: To prepare, combine 45 mL of Milli-Q® and 5 mL of 10X PBS.
2. 1X PBST for washes: To prepare, combine 45 mL of Milli-Q®, 5 mL of 10X PBS, and lastly, 50 µL of Tween 20 (0.1%).
3. 5X SSCT for washes: To prepare, combine 30 mL of Milli-Q®, 10 mL of 20X SSC, and lastly, 40 µL of Tween 20 (0.1%).
4. Proteinase K solution for permeabilization: To prepare 4 mL of a 10 µg/ml solution, combine 4 mL of 1X PBS and 4 µL of 10 mg/ml proteinase K.
5. Bleaching solution (optional-FFPE protocol): To prepare 1 mL, combine in this order: 962.3 µL H_2_O, 20µL H_2_O_2_ and 17.7 µL KOH.
6. 1X Tris-EDTA Buffer (FFPE protocol): For 50 mL, combine 49.5 mL of 1X PBS and 0.5mL of 100X TRIS EDTA buffer pH 9.0.

## 3. Detailed Methods

Both protocols are performed over three consecutive days, excluding tissue processing, sectioning, and imaging. Detailed step-by-step procedures for both workflows are provided in Supplementary File 1 and Supplementary File 2.

### 3.1 In silico workflow for Probe design (optional)

As an optional component of this study, we built a step-to-step workflow to assist with in silico probe design prior to experimental validation because genome annotation in *P. waltl* remains incomplete. The workflow can be divided into three main stages that guide the identification of suitable transcript regions for HCR probe generation: (i) target transcript selection, (ii) orthology confirmation, and (iii) probe design for downstream procedures as oligonucleotide synthesis (Step 5) and experimental detection (Step 6). A schematic overview is provided in Figure 2, and full step-by-step instructions are included in Supplementary File 3.

**Figure 2.**
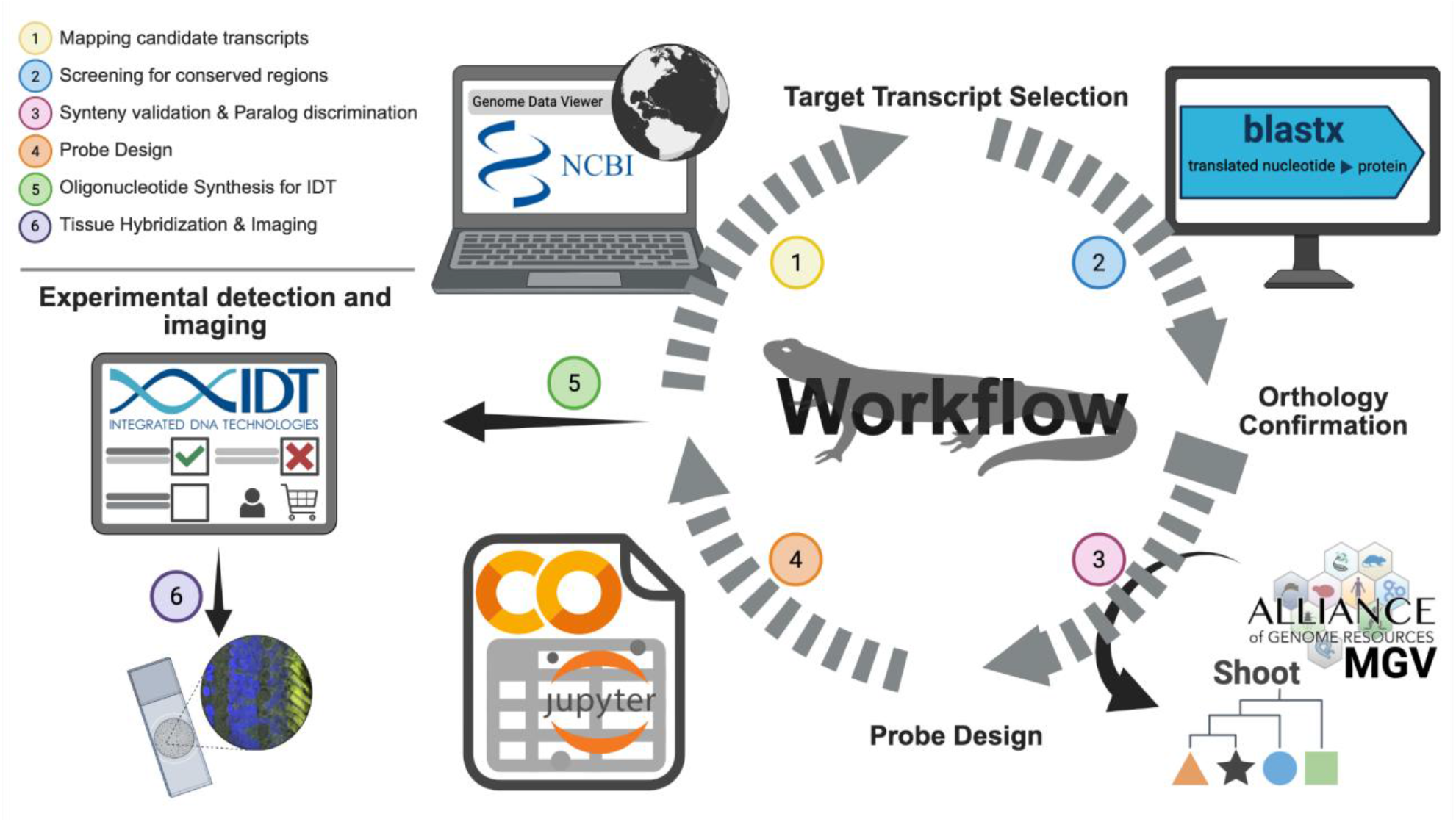
Probe design in silico to *i*n situ HCR-FISH detection in *P. waltl*. Schematic overview of the workflow linking: (**Steps 1-2**) Target transcript selection by mapping candidates from NCBI and screening conserved regions; (**Step 3**) Orthology confirmation by synteny analysis and paralog discrimination to define the sequence target; (**Step 4**) Probe design implemented as a Google Colab notebook to generate split-initiator probe sets for synthesis; (**Step 5**) low-cost oligonucleotide synthesis through IDT to enable multiplex targeting; and (**Step 6**) experimental detection by in situ HCR-FISH and imaging. *Created in BioRender. Bendezu, S. (2026) https://BioRender.com/ivwaq9s*

In Step 1, candidate transcripts were selected using NCBI Genome Data Viewer (Rangwala et al. 2021). In Step 2, conserved regions were screened using BLASTX (Johnson et al. 2008) against human and *Xenopus tropicalis*, together with axolotl and *Gallus gallus* to improve its detection. In Step 3, orthology was further assessed by combining synteny visualization in Multiple Genome Viewer (Richardson et al. 2022) with phylogenetic placement using SHOOT (Emms & Kelly 2022) to support orthology confirmation and help discriminate potential paralogs. In Step 4, probe sets were designed using a Google Colab-based implementation that leverages cloud computing and builds on the open-source HCR 3.0 Probe Maker, originally developed by the Özpolat Lab under the GNU General Public License 3.0 (Choi et al. 2018); (Kuehn et al. 2022); (Tsuneoka & Funato 2020); (Wang et al. 2024). In Step 5, selected probe sets were exported as an order-ready Excel file for oligonucleotide synthesis to Integrated DNA Technologies (IDT). Finally, Step 6 is followed by experimental detection and imaging through HCR-FISH in *P. waltl*.

Users can upload organism-specific cDNA datasets to generate candidate probe sequences using the Google Colab implementation. In Section III (Upload Data) of the Colab_Probe_Tool.ipynb notebook, FASTA files can be selected from the existing folder or uploaded directly for candidate probe generation. A more detailed description of this step is provided in Suppl. File 3 (Step 5, Section 3.1.3, “Probe design for IDT ordering”).

In addition to *P. waltl*, the probe tool has been used with *Gallus gallus* transcriptomic dataset, demonstrating its utility for both emerging and established model organisms. Final probe sets were synthesized by Integrated DNA Technologies and experimentally validated against Molecular Instruments designed probes using HCR-FISH in *P. waltl* ocular tissue. The Google Colab implementation (google.colab 0.0.1a2) performed in this study uses Python 3.12.12, and the Colab notebook is available through our GitHub repository. We set up the tool to improve its accessibility and usability for the broader community. See the “Code Availability” section for details.

### 3.2 Tissue collection and fixation (Day 0)

Larval or adult *P. waltl* eye tissues were collected and immediately fixed to preserve RNA integrity. Two tissue processing workflows were used in this study depending on the downstream imaging strategy: whole-mount preparation followed by cryosectioning and formalin-fixed paraffin-embedded (FFPE) tissue protocol.

For whole-mount experiments, samples were fixed in 4% paraformaldehyde (PFA) for 1h at room temperature and then transferred to 100% methanol for dehydration and storage. For paraffin processing, embryos were fixed in 10% buffered formalin for 1h at 4ºC before standard paraffin embedding and sectioning.

Fixation time was optimized for *P. waltl* tissue. Initial experiments followed the Molecular Instruments HCR-FISH v3.0 system protocol developed for chicken embryos, which recommends fixation for up to 24 hours. However, reducing fixation time to one hour resulted in improved signal intensity (Fig. 3a and 3b). Because PFA fixation promotes protein and nucleic acid cross-linking, this may explain reduced probe accessibility to target RNA transcripts (Kiernan 2000); (Hoffman et al. 2015). A shorter fixation period preserved tissue morphology while improving probe penetration and signal detection in newt eye tissues.

**Figure 3.**
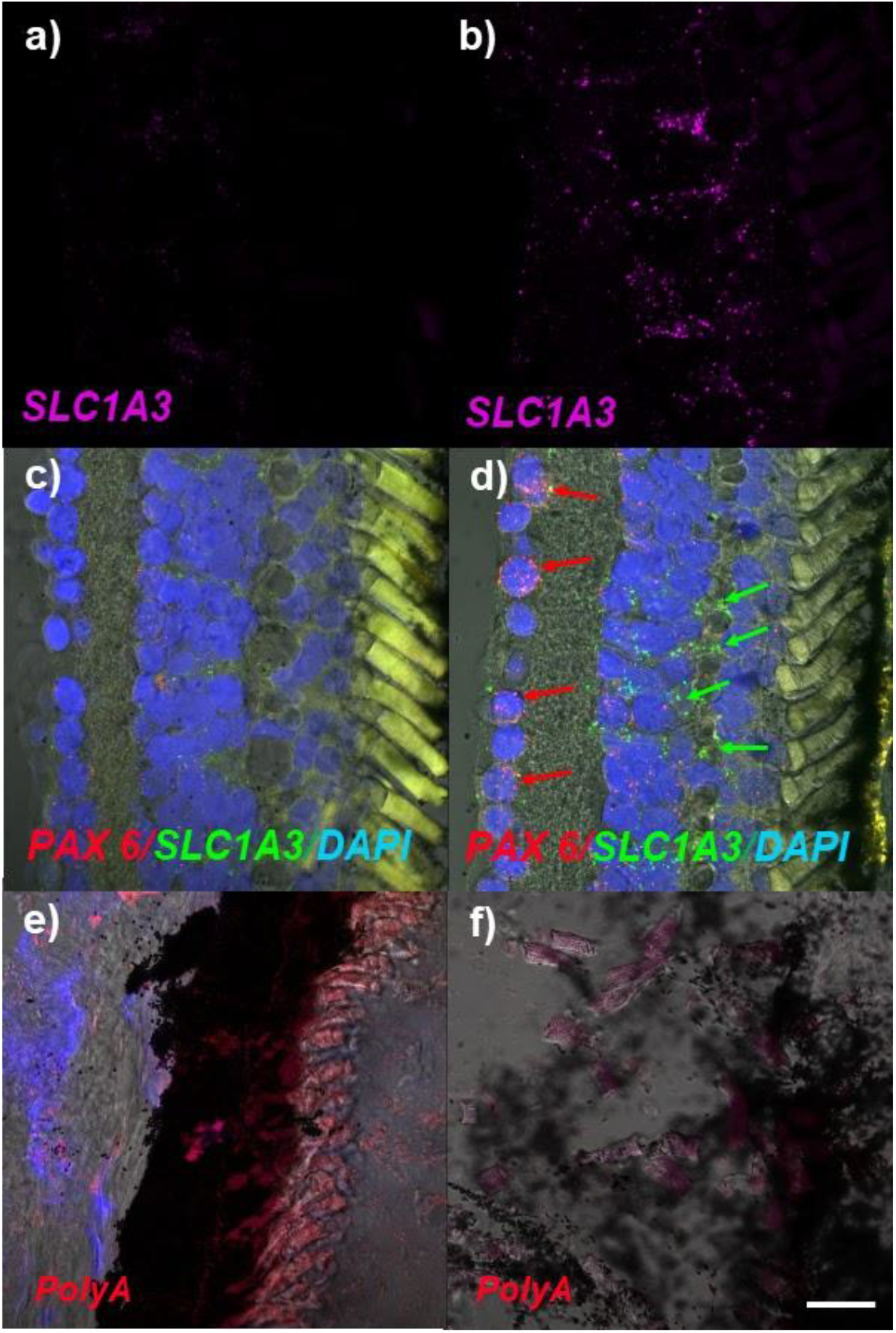
Manipulation of variables in the HCR-FISH whole-mount protocol followed by cryosection. (**a**) Section of newt retina with *SLC1A3*, a marker for Müller glial cells, after PFA fixation overnight. (**b**) *SLC1A3* expression in newt retina after one-hour fixation with PFA. (**c)** *PAX6* and *SLC1A3* expression within the retina after 100 µg/mL proteinase K treatment for 3 minutes. (**d**) *PAX6* and SLC1A3 expression (indicated by arrows) within the retina after 10 µg/mL proteinase K treatment for 3 minutes. (**e**) Poly A, a positive control for mRNA transcripts, expression in a retina treated with 10 µg/mL proteinase K for 45 minutes. (**f**) No visible Poly A signal within the retina after treatment with 30 µg/mL proteinase K for 45 minutes. The scale bar in f is 20 µm and applies to all panels.

### 3.3 Tissue preparation and Probe Hybridization (Day 0 and 1)

#### 3.3.1 Whole-mount workflow

For the whole-mount workflow, methanol-stored samples were first rehydrated through a graded methanol series in PBST and then washed in PBST prior to permeabilization. Samples were treated with proteinase K to improve probe penetration, followed by a brief post-fixation in 4% PFA to stabilize tissue morphology after digestion. For highly pigmented tissues, a brief bleaching step using hydrogen peroxide and potassium hydroxide was optionally performed under strong illumination to reduce pigmentation and improve visualization of fluorescent signals. After additional washes in PBST and SSCT, samples were equilibrated in probe hybridization buffer before incubation with probe solution overnight at 37 °C. Detailed step-by-step procedures are provided in Supplementary File 1.

#### 3.3.2 FFPE workflow

For the FFPE workflow, paraffin sections mounted on Superfrost™ Plus slides were first baked at 60 °C to improve tissue adherence and then deparaffinized using sequential xylene and ethanol washes. Slides were subsequently rehydrated through a graded ethanol series before antigen retrieval was performed using a heated Tris-EDTA buffer. After cooling, slides were washed in PBST and prepared for hybridization by creating a hydrophobic barrier around the tissue sections with a Peroxidase-Antiperoxidase (PAP) pen.

In some experiments, additional permeabilization was performed using proteinase K digestion prior to hybridization. This step was included to improve probe penetration in thicker sections but was not required for all samples. For highly pigmented tissues, a brief bleaching step using hydrogen peroxide and potassium hydroxide was optionally performed under strong illumination to reduce pigmentation and improve visualization of fluorescent signals.

Following permeabilization and optional bleaching, slides were equilibrated in probe hybridization buffer prior to probe incubation. Probe sets were diluted in probe hybridization buffer according to manufacturer recommendations, with concentrations ranging from 4–16 nM depending on transcript abundance. For initial probe testing, a concentration of 16 nM was used, which represents the maximum recommended concentration for split-initiator probes. In our experiments, probe solutions were typically prepared by adding 1.6 µL of each probe set from a 1 µM stock to 100 µL of probe hybridization buffer. Approximately 100 µL of probe solution was applied to each slide, and samples were incubated overnight at 37 °C in a humidified chamber. Detailed step-by-step procedures for the FFPE workflow are provided in Supplementary File 2.

#### 3.3.3 Proteinase K Optimization

Proteinase K digestion was optimized to balance tissue permeability with preservation of structural integrity. Proteinase K facilitates probe access by digesting proteins within the tissue matrix, including nucleases that may degrade RNA. The Molecular Instruments HCR-FISH v3.0 system protocol for chick embryos recommends a concentration of 10 µg/mL, whereas the zebrafish larvae protocol suggests 30 µg/mL. To determine the optimal conditions for *P. waltl*, we tested proteinase K at concentrations of 10 µg/mL, 30 µg/mL, and 100 µg/mL. Higher concentrations resulted in significant tissue degradation by the end of the protocol (Fig. 3c and 3d).

We also evaluated incubation duration. The Molecular Instruments zebrafish protocol recommends digestion for up to 45 minutes, while the chick embryo protocol suggests a maximum of 2.5 minutes. We conducted an experiment to test the 45-minute incubation at two concentrations: 10 µg/mL and 30 µg/mL (Figure 3e and 3f). Extended incubations produced severe degradation regardless of concentration (Figure not shown). Based on these results, a concentration of 10 µg/mL for 3 minutes was selected as the optimal condition for *P. waltl* samples, preserving tissue integrity while allowing adequate probe penetration.

### 3.4 HCR Probe Amplification (Day 2)

#### 3.4.1 Whole-mount workflow

Following probe hybridization, whole-mount samples were rinsed in a pre-warmed probe wash buffer at 37 °C to remove excess probes, followed by additional washes in SSCT at room temperature. Hairpin amplifiers (H1 and H2) were denatured by brief heating and allowed to cool to room temperature in the dark to enable proper folding. Samples were then equilibrated in amplification buffer prior to incubation with the hairpin solution. Amplification was carried out overnight at room temperature in the dark to allow signal development. All steps were performed with gentle agitation to preserve tissue integrity.

#### 3.4.2 Paraffin workflow

Following probe hybridization, paraffin sections were washed through a graded series of probe wash buffer and SSCT at 37 °C to remove excess probe and reduce background signal, followed by a final wash in SSCT at room temperature. Slides were then equilibrated in amplification buffer prior to signal amplification. Hairpin amplifiers (H1 and H2) were denatured by brief heating and allowed to cool in the dark to ensure proper folding before use. Amplification buffer containing hairpins was applied directly to tissue sections, and slides were incubated overnight at room temperature in a dark, humidified chamber to allow signal development. Care was taken throughout to maintain hydration of tissue sections and to minimize light exposure.

### 3.5 Probe washing, Image preparation, and Sectioning (Day 3)

#### 3.5.1 Whole-mount workflow

Following HCR amplification, excess hairpins were removed by washing samples in 5X SSCT buffer at room temperature in the dark with gentle rocking. Samples designated for whole-mount imaging may be left in SSCT for several days and a protocol for optical clearing can be performed after this step.

On the other hand, samples designated for cryosectioning were subsequently transferred to 30% sucrose and incubated at 4 °C until the tissue sank, allowing cryoprotection and dehydration prior to embedding. Samples were then embedded in OCT compound, stored at −80 °C, and protected from light until sectioning.

Cryosections were prepared at the desired thickness using a cryostat in a low-light environment to preserve fluorescence. Sections were mounted on Superfrost™ Plus slides and stored at −20 °C until further processing. Prior to imaging, slides were washed briefly in PBST followed by PBS. Nuclear staining was performed using DAPI, after which slides were washed in PBS and mounted with Fluoromount™ before coverslipping.

#### 3.5.2 FFPE workflow

Following overnight amplification, FFPE sections were washed in 5X SSCT at room temperature in the dark to remove excess hairpins while preserving fluorescence. Slides were then counterstained with DAPI, washed in PBS, and mounted with Fluoromount™ prior to coverslipping. To minimize photobleaching, samples were protected from light throughout the post-amplification washing and mounting steps.

## 4. Results

After much trial and error, we were able to optimize the HCR-FISH protocol in our emerging model organism, the Iberian ribbed newt, by manipulating various variables throughout the trials. Starting with fixation time, we decided on fixing our whole-mount samples for an hour before transferring to methanol because we saw more robust signal with less fixation (Figure 3a and b). With the proteinase K solution and incubation, we manipulated the concentration and duration of exposure to this solution. We found that a lower concentration of proteinase K resulted in a more robust signal, and a shorter digestion time preserved the integrity of the tissue better (Figures 3c-f). We validated probe markers in the newt retina including *SLC1A3* and *PAX6*, each marking for Müller glial and ganglion cells respectively (Figure 3a-d). To validate our in silico probe design (Figure 2), we used an *RPE65* probe targeting retinal pigment epithelium cells and compared its signal to that obtained with a probe designed by Molecular Instruments. The two probes exhibited comparable signal intensity and spatial localization, confirming the reliability of probe design using this extended in silico workflow (Figure 4). We further validated this approach in FFPE retinal sections using the same RPE65 probes and again observed comparable signal intensity and spatial localization relative to the Molecular Instruments probe (Figure 5).

**Figure 4.**
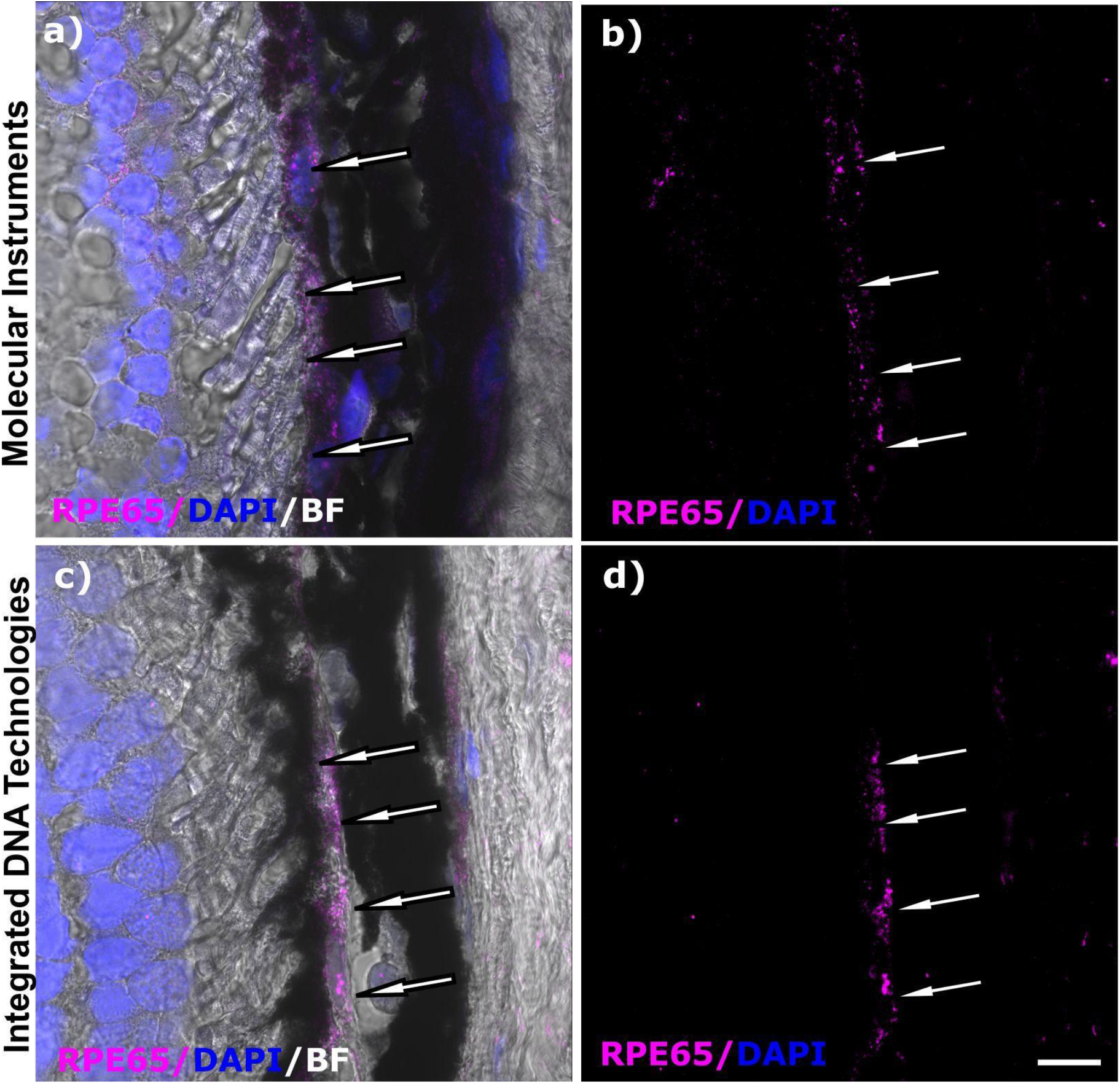
Validation of the *RPE65* HCR-FISH (whole-mount protocol followed by cryosectioning) probe using our step-to-step in silico workflow. The *RPE65* probe was designed via our cloud-deployed Google Colab implementation. This probe marks retinal pigment epithelium cells (indicated by the arrows). DAPI was used to counter stain the nuclei. The scale bar in d is 20µm and applies to all the panels. Note: A brief 3-minute bleaching step was performed with these samples according to our optimized protocol in Supplementary Material 1.

**Figure 5.**
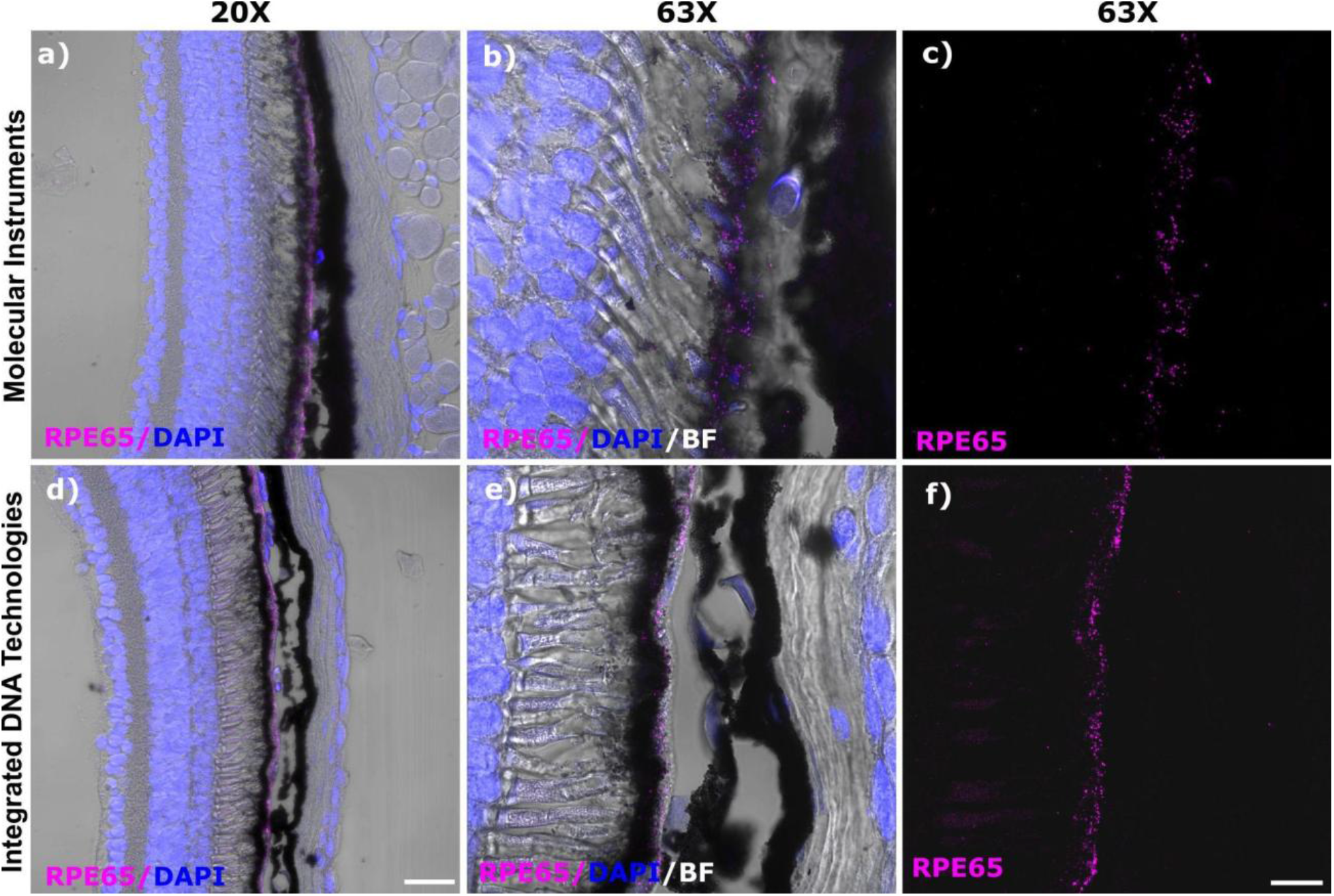
Validation of the RPE65 HCR-FISH (FFPE protocol) probe using our step-to-step in silico workflow. The *RPE65* probe was designed via our cloud-deployed Google Colab implementation.This probe marks retinal pigment epithelium cells. DAPI was used to counter stain the nuclei. The scale bar in a is 50µm and applies to **d**. The scale bar in **f** is 20µm and applies to panels **b**,**c** and **e**. Note: A brief 30-second bleaching step was performed according to our optimized protocol in Supplementary File 2.

## 5. Potential Pitfalls/Troubleshooting

We found that there was potential for error with the fixation of tissues in preparation for the HCR protocol. We determined that prolonged fixation reduces signal intensity, likely due to increased cross-linking that limits probe accessibility (Kiernan 2000); (Hoffman et al. 2015). A one-hour fixation was optimal for whole-mount eyes but we recommend optimizing based on individual tissue type.

Another area for potential pitfalls involves the overdigestion of tissues during the probe hybridization step. Proteinase K treatment has been used for ISH and in situ HCR to improve the penetration of RNA/DNA probes and fluorophore-labeled hairpins by digesting proteins (Tsuneoka & Funato 2020). However, proteinase K treatment could damage the morphology of the tissue and the integrity of some proteins. When tested, higher concentrations (≥30 µg/mL of proteinase K) or prolonged incubation (≥45 minutes) resulted in tissue degradation and diminished signal. A concentration of 10 µg/mL for 3 minutes preserved tissue integrity while maintaining probe penetration in whole-mounts. In some cases for the FFPE protocol, no proteinase K treatment resulted in better preserved tissue sections and a robust hybridization signal, highlighting the potential need to troubleshoot this step.

Lastly, imaging intact pigmented whole-mount eyes can reduce fluorescent signal visibility. Cryosecting after the whole-mount protocol resolved this limitation without compromising signal localization. A brief optional bleaching step was integrated into both the whole-mount and FFPE protocol using hydrogen peroxide and potassium hydroxide to circumvent these pigmentation-related issues. Additionally, optical clearing after whole-mount processing, as it has been performed in *P. waltl* brain (Woych et al. 2022), can be conducted to improve and optimize the 3D-visualization for eyes that inherently have pigmented tissues.

## 6. Discussion/Future Directions

Future work will focus on expanding the set of experimentally validated probes generated through the in silico workflow described in Section 3.1 and applying this pipeline to additional markers relevant to retina and lens development and regeneration. Broader probe validation will strengthen the utility of HCR-FISH as a complement to RNA sequencing datasets in *P. waltl* and support spatial confirmation of candidate transcripts identified through transcriptomic analyses.

In the long-term, our results would be further strengthened by quantifying the number of RNA molecules expressed for our genes of interest within the eye. This would allow us, for example, to measure gene expression changes that are taking place in the regeneration response to injury over time. Because multiplex imaging in this system remains constrained by tissue pigmentation and signal visualization, continued optimization of bleaching and tissue-clearing conditions for whole-mount visualization (Woych et al. 2022) will be important, particularly for preserving tissue morphology and fluorescence in highly pigmented ocular tissues.

Together, these efforts should further improve the use of HCR-FISH in *P. waltl* and support future studies aimed at mapping gene expression dynamics during eye development and regeneration.

## Code availability

The notebook used for the Probe design (4 step) and its supporting scripts are available in the GitHub repository at https://github.com/DelRioTsonisLab/HCR-probes-Tool The repository includes detailed instructions for running the workflow in Google Colab or locally, a dependency list (packages.txt), and a worked example (Vignette.html) for users running the tool locally.

## Acknowledgements/Funding Sources

This work was supported by funding sources at Miami University to KDRT, including from the Department of Biology and the National Institute of Health grant EY035325. SMR was supported by various funding sources at Miami University including the Louis Stokes Alliance for Minority Participation (LSAMP), Undergraduate Research Award (URA), and Dean’s Scholar Award. SMR was also supported by SDB “Choose Development!” fellowship. We thank Dr. Nicholas Leigh from Lund University and his lab for hosting SMR and generously lending insights to help optimize this protocol. The authors further thank Dr. Byran Smucker on experimental design advise, and Dr. Zachery Oestreicher, the Director of the Center for Advanced Microscopy and Imaging (CAMI) at Miami University for instrumentation support.

## Ethics Approval

Handling and surgical procedures were performed following guidelines by the Institutional Animal Care and Use Committee at Miami University.

## Supplementary material

Supplementary File 1 provides a step-by-step user protocol for the optimized HCR-FISH Protocol in whole-mount and fixed frozen *Pleurodeles waltl* tissue.

Supplementary File 2 provides a step-by-step user protocol for the optimized HCR-FISH Protocol in Formalin-Fixed Paraffin-Embedded (FFPE) *Pleurodeles waltl* tissue.

Supplementary File S3 provides a step-by-step user guide (manual) for the optional in-silico workflow for Probe design used in this study.

## References

Brown, T. et al., 2025. Chromosome-scale genome assembly reveals how repeat elements shape non-coding RNA landscapes active during newt limb regeneration. Cell Genomics, 5(2), p.100761. Available at: 10.1016/j.xgen.2025.100761.

Choe, K. et al., 2023. Advances and challenges in spatial transcriptomics for developmental biology. Biomolecules, 13(1), p.156. Available at: 10.3390/biom13010156.

Choi, H.M.T. et al., 2018. Third-generation in situ hybridization chain reaction: multiplexed, quantitative, sensitive, versatile, robust. Development (Cambridge, England), 145(12), p.dev165753. Available at: 10.1242/dev.165753.

Ding, J. et al., 2020. Systematic comparison of single-cell and single-nucleus RNA-sequencing methods. Nature Biotechnology, 38(6), pp.737–746. Available at: 10.1038/s41587-020-0465-8.

Elewa, A. et al., 2017. Reading and editing the Pleurodeles waltl genome reveals novel features of tetrapod regeneration. Nature Communications, 8(1), p.2286. Available at: 10.1038/s41467-017-01964-9.

Emms, D.M. & Kelly, S., 2022. SHOOT: phylogenetic gene search and ortholog inference. Genome Biology, 23(1), p.85. Available at: 10.1186/s13059-022-02652-8.

Garza-Garcia, A.A., Driscoll, P.C. & Brockes, J.P., 2010. Evidence for the local evolution of mechanisms underlying limb regeneration in salamanders. Integrative and Comparative Biology, 50(4), pp.528–535. Available at: 10.1093/icb/icq022.

Hoffman, E.A. et al., 2015. Formaldehyde crosslinking: a tool for the study of chromatin complexes. The Journal of Biological Chemistry, 290(44), pp.26404–26411. Available at: 10.1074/jbc.R115.651679.

Huang, T. et al., 2023. A rapid and sensitive, multiplex, whole mount RNA fluorescence in situ hybridization and immunohistochemistry protocol. Plant Methods, 19(1), p.131. Available at: 10.1186/s13007-023-01108-9.

Jin, L. & Lloyd, R.V., 1997. In situ hybridization: Methods and applications. Journal of Clinical Laboratory Analysis, 11(1), pp.2–9. Available at: 10.1002/(SICI)1098-2825(1997)11:1<2::AID-JCLA2>3.0.CO;2-F.

Johnson, M. et al., 2008. NCBI BLAST: a better web interface. Nucleic Acids Research, 36(Web Server issue), pp.W5–9. Available at: 10.1093/nar/gkn201.

Kiernan, J.A., 2000. Formaldehyde, formalin, paraformaldehyde and glutaraldehyde: What they are and what they do. Microscopy Today, 8(1), pp.8–13. Available at: 10.1017/s1551929500057060.

Kirkham, M. & Joven, A., 2015. Studying newt brain regeneration following subtype specific neuronal ablation. Methods in Molecular Biology (Clifton, N.J.), 1290, pp.91–99. Available at: 10.1007/978-1-4939-2495-0_7.

Kuehn, E. et al., 2022. Segment number threshold determines juvenile onset of germline cluster expansion in Platynereis dumerilii. Journal of Experimental Zoology. Part B, Molecular and Developmental Evolution, 338(4), pp.225–240. Available at: 10.1002/jez.b.23100.

Matsunami, M. et al., 2019. A comprehensive reference transcriptome resource for the Iberian ribbed newt Pleurodeles waltl, an emerging model for developmental and regeneration biology. DNA Research: An International Journal for Rapid Publication of Reports on Genes and Genomes, 26(3), pp.217–229. Available at: 10.1093/dnares/dsz003.

National Eye Institute (NEI), 2016. Visual impairment, blindness cases in U.S. expected to double by 2050. Available at: https://www.nei.nih.gov/about/news-and-events/news/visual-impairment-blindness-cases-us-expected-double-2050 [Accessed April 8, 2026].

Rangwala, S.H. et al., 2021. Accessing NCBI data using the NCBI Sequence Viewer and Genome Data Viewer (GDV). Genome Research, 31(1), pp.159–169. Available at: 10.1101/gr.266932.120.

Richardson, J.E., Baldarelli, R.M. & Bult, C.J., 2022. Multiple genome viewer (MGV): a new tool for visualization and comparison of multiple annotated genomes. Mammalian Genome: Official Journal of the International Mammalian Genome Society, 33(1), pp.44–54. Available at: 10.1007/s00335-021-09904-1.

Tsonis, P.A. & Del Rio-Tsonis, K., 2004. Lens and retina regeneration: transdifferentiation, stem cells and clinical applications. Experimental Eye Research, 78(2), pp.161–172. Available at: 10.1016/j.exer.2003.10.022.

Tsuneoka, Y. & Funato, H., 2020. Modified in situ hybridization chain reaction using short hairpin DNAs. Frontiers in Molecular Neuroscience, 13, p.75. Available at: 10.3389/fnmol.2020.00075.

Wang, Y. et al., 2024. Multiplexed in situ protein imaging using DNA-barcoded antibodies with extended hybridization chain reactions. Nucleic Acids Research, 52(15), p.e71. Available at: 10.1093/nar/gkae592.

Woych, J. et al., 2022. Cell-type profiling in salamanders identifies innovations in vertebrate forebrain evolution. Science (New York, N.Y.), 377(6610), p.eabp9186. Available at: 10.1126/science.abp9186.

Zhang, F. et al., 2002. Differential regulation of fibroblast growth factor receptors in the regenerating amphibian spinal cord in vivo. Neuroscience, 114(4), pp.837–848. Available at: 10.1016/s0306-4522(02)00321-4.

