## Supplementary file 1 for "Optimizing the hybridization chain reaction-fluorescence in situ hybridization (HCR-FISH) protocol for *Pleurodeles waltl*"

**Supplementary File 1**
**Optimized HCR-FISH Protocol for whole-mount and fixed frozen *Pleurodeles waltl* tissue***

*Use Molecular Grade reagents throughout. Use standard practices for maintaining an RNase free environment.*

Day 0: Specimen collection (Duration: 1 hour + Overnight (ON))

1. Collect larvae or wholemount organ in 4% PFA for 1 hour.
2. Transfer to 100% methanol overnight and store for up to 6 months.

Day 1 HCR: Permeabilization and Hybridization (Duration: 3 hours + ON)

1. Thaw probe hybridization buffer (stored at -20°C) at room temperature. Place PBST, SSCT, and probe sets on ice to chill.
    a. Warning: Probe hybridization buffer is toxic (use gloves and work under the hood).
2. Rehydrate samples in a graded methanol wash, each for 5 minutes at room temperature:
    a. 75% methanol / PBST
    b. 50% methanol / PBST
    c. 25% methanol / PBST
3. Wash samples three times at room temperature with PBST. Prepare proteinase K solution during washes.
4. Treat samples with proteinase K solution for 3 minutes at room temperature
5. Wash samples 3x with PBST at room temperature. Thaw 4% PFA during washes.
6. Post-fix sample for 20 minutes in 4% PFA at room temperature.
7. Wash samples 5x with 1X PBST on ice.
   *Steps 8 to 10 optional for highly pigmented organs or tissue*:
8. Bleach samples in bleaching solution at room temperature under a bright light for 5 minutes, turning the sample often to assure an even bleaching process. Time has to be optimized depending on how pigmented the tissue is and the size.
9. Wash samples 3x with PBS at room temperature.
10. Degas samples under a vacuum for 15 minutes on ice. This step can be performed by placing the specimen on PBS in the bottom of a 50ml Corning filtration tube used for bacterial filtration. Place the tube in an ice bucket on a shaker and attach it to a vacuum.
11. Wash samples 2x with 5X SSCT on ice.
12. Transfer larvae or organ(s) to a 2 mL Eppendorf tube. Pre-hybridize samples with 250-500 µl of probe hybridization buffer for 5 minutes at room temperature.
13. Remove the buffer, then repeat with 500 µl of probe hybridization buffer for 30 minutes at 37 °C. While incubating, proceed to the next step.

14. Prepare probe solution by adding 16 nM (8 μL) of each initiator probe set to 500 μl of probe hybridization buffer. Pre-warm solution to 37°C. (Ratio: 1.6 μL per 100 μL of probe hybridization buffer).
15. Remove the pre-hybridization solution and add the prepared probe solution.
16. Incubate the samples overnight at 37ºC.
    a. Optional: Parafilm seal the tube to prevent evaporation.

<u>Day 2 HCR: Probe wash and Amplification (Duration: 3 hours + ON, little hands-on time)</u>

1. Thaw amplification buffer at room temperature and wash buffer at 37°C. Turn on the heating block to 95°C. Wait ~ 1 hour for all solutions to equilibrate.
    a. Buffers are toxic (use gloves and work under the hood).
2. Remove probe solution carefully with pipette. Wash samples 4 X 15 min with 1 mL of pre-warmed probe wash buffer at 37 ºC, with gentle rocking. During the third wash, proceed to the next step.
3. Briefly spin down hairpin aliquots and heat at 95ºC for 90 seconds. Cool aliquots to room temperature in a dark drawer for 30 min. Turn off the heating block. While cooling, proceed to the next step.
4. Wash samples 2x 5 min with 5 mL of 5X SSCT at room temperature.
5. Transfer embryos to a clean 1.5 mL Eppendorf tube. Pre-amplify samples with 500 μL of amplification buffer for 5 minutes at room temperature.
6. In the dark, add 10 μL each of cooled hairpins H1 and H2 to 480 μL of room temperature amplification buffer. **[Subsequent steps should be performed in the dark]**
7. Remove the pre-amplification solution and add the hairpin solution.
8. Incubate the samples overnight in the dark at room temperature.
    a. Optional: Parafilm seal the tube.

<u>Day 3: Hairpin washing and Optional Cryoembedding (Duration: 3 hours 15 min)</u>

1. Remove excess hairpins by washing with 500 μl of 5X SSCT at room temperature, with gentle rocking, in the dark (Tip: Cover slides with aluminum foil to protect from light).
    a. 2 X 5 min
    b. 2 X 30 min
    c. 1 X 5 min
2. NOTE: If you are planning to image wholemount, you may leave the sample in SSCT for several days or follow a protocol for optical clearing. If you plan to section, proceed to steps 3 and 4, and sectioning section.
3. Transfer to 10 mL of 30% sucrose for at least two hours or until the samples sink to allow for the sucrose to penetrate and dehydrate the tissue. Perform this step in the dark at 4 ºC.

4. On the same day, samples left in sucrose should be cryo-embedded in OCT, stored at - 80 ºC, and protected from light before sectioning.

Optional: Sectioning (Duration: 90 minutes)

1. Cryosection at appropriate thickness in a low-light environment. Keep sections at -20 °C until ready to coverslip. Use Fisherbrand™ Superfrost™ Plus Microscope Slides for maximum adherence.
2. On same day, treat slides as follows:
    a. 5 mins PBST
    b. 5 mins PBS
    c. 60 mins DAPI
    d. 2 x 5 mins PBS
    e. Coverslip w/ Fluoromount™

*Adapted from Molecular Instruments HCR-FISH v3.0 system Protocols for Whole-mount Chicken Embryo and Zebrafish Larvae. https://www.molecularinstruments.com/
