## Supplementary File 2 for "Optimizing the hybridization chain reaction-fluorescence in situ hybridization (HCR-FISH) protocol for *Pleurodeles waltl*"

**Supplementary File 2**
**Optimized HCR-FISH Protocol for FFPE *Pleurodeles waltl* tissue***

*Use Molecular Grade reagents throughout. Use standard practices for maintaining an RNase free environment.*

Day 0: Specimen collection, Tissue Processing, and Sectioning (Duration: 1 hour + Overnight (ON))

1. Collect embryos into molecular-grade PBS. Rinse with PBS several times, if necessary.
2. Place embryos into 10% buffered formalin for 1 hour to fix at 4°C.
3. Wash 3x with molecular grade PBS.
4. For E7 chickens and older, make a small incision through the anterior chamber of the eye to vent. This will prevent collapse of the eye during paraffinization.
5. Dissect off any excess tissue, as necessary.
6. Place samples in 70% EtOH at 4°C for short-term (30 min to overnight) storage.
7. Run the tissue processor using standard protocol for paraffin embedding and embed.
8. Section paraffin blocks according to standard protocol using Fisherbrand™ Superfrost™ Plus Microscope Slides for maximum adherence.

Day 1 HCR: Permeabilization and Hybridization (Duration: ~3 hours + ON)

1. Thaw probe hybridization buffer (stored at -20°C) at room temperature.
2. Bake slides in an oven at 60 °C for one hour. (Tip: Label all slides with pencil, don't use sharpie as it will come off in ethanol washes.)
3. Deparaffinize samples by placing them in a Coplin jar with fresh xylenes 3x 5 minutes.
4. Incubate slides in 100% EtOH for 3x 3 minutes at room temperature.
5. Rehydrate samples in a graded ethanol wash, each for 3 minutes at room temperature (prepare fresh solutions):
    a. 95% EtOH in 1X PBS
    b. 70% EtOH in 1X PBS
    c. 50% EtOH in 1X PBS
    d. 1X PBS
6. Prepare 50 mL of the 1X Tris-EDTA buffer in a Coplin jar, using PBS to dilute. Heat in the microwave until boiling (watch carefully).
7. Immediately place slides in the hot Tris-EDTA. Place the slides in the 60°C oven to maintain a high temperature. Incubate for 15 minutes.
8. Remove slides from the jar and cool for 5 minutes in water, replace the water and let the slides sit for 5 more minutes. While waiting, place a humidified chamber at

37°C to preheat for later steps. Place probe sets on ice to thaw.

9. Immerse slides in 1X PBST in a Coplin jar for 2x 2 minutes. Prepare fresh proteinase K solution during washes and/or bleaching solutions (optional).
10. Dry the edges of the slides with a Kimwipe™ without touching the tissue. Create a barrier with a hydrophobic pen.
11. Optional: Add 200 µl proteinase K solution to each slide. Incubate 10 minutes at 37°C in a humidified chamber.
12. For pigmented samples only: Bleach slides in a bleaching solution at room temperature under a bright gooseneck lamp for 30 seconds (optimize the time for the thickness of your sample)
13. Wash slides 2x in fresh PBST in a Coplin jar for 3 minutes each at room temperature. Return the humidified chamber to 37°C.
14. Add 200 µl of probe hybridization buffer to each slide. Pre-hybridize for 10 minutes in a humidified chamber.
15. While incubating, prepare the probe solution. For a 16nM concentration, add 1.6 µl of each initiator probe set for each 100 µl of probe hybridization buffer.
16. Gently pour the buffer from pre-hybridization off slides into a formamide waste container. Without letting the slides dry, add 100 µl of probe solution to each slide. Add probe hybridization buffer to negative control slides.
17. Using forceps, carefully position a piece of broken coverslip on the top and bottom of the slide to create a platform. Place a full coverslip over each sample. The broken coverslip shards should prevent the coverslip from laying directly on the sections and will facilitate easy removal later. Try to position the coverslip so that there are no bubbles on top of sections. Make sure to not cross-contaminate probe sets with the forceps.
    a. Tip: This step can be skipped if a beaker with water is placed in the oven to make sure the chamber is humid.
18. Incubate slides overnight at 37°C in a humidified chamber.

<u>Day 2 HCR: Probe wash + Amplification (Duration: ~3 hours + ON, little hands-on time)</u>

1. Thaw amplification buffer at room temperature and place probe wash buffer at 37ºC. Turn on the heating block to 95°C. Wait ~ 1 hour for all solutions to equilibrate.
2. Carefully remove coverslips and collect broken shards into glass waste.
3. Wash slides by adding 200 µl of the following solutions in sequence for 15 minutes each at 37°C. Collect all waste into a formamide waste container and use a humidified chamber during incubations:
    a. 75% probe wash buffer / 5X SSCT
    b. 50% probe wash buffer / 5X SSCT

c. 25% probe wash buffer / 5X SSCT
d. 100% 5X SSCT [Proceed to step 8 during this incubation period]

4. Immediately place slides in a Coplin jar containing fresh 5X SSCT for 5 minutes at room temperature.
5. Dry the edges of slides with a Kimwipe™. Re-apply barrier with hydrophobic pen, if necessary.
6. Pipette 200 µl of amplification buffer on each slide and pre-amplify in a humidified chamber for 30 minutes at room temperature. Proceed to the next step during incubation.
7. Briefly spin down hairpin aliquots and heat at 95ºC for 90 seconds. Cool aliquots to room temperature in a dark drawer for 30 min. Turn off the heating block. While cooling, proceed to the next step.
8. In the dark, add 1 µl of each hairpin for each 100 µl amplification buffer at room temperature. **[Subsequent steps should be performed in the dark]**
9. Drain the pre-amplification solution from slides into a formamide waste container. Blot the edges of the slide dry with a Kimwipe™. Without letting the sections dry, add 100 µl of the hairpin solution to each slide.
10. Incubate slides overnight in a dark, humidified chamber at room temperature.

<u>Day 3: Hairpin washing (Duration: 1 hour)</u>

1. Remove excess hairpins on slides by washing with 200 µl of 5X SSCT at room temperature in the dark (Tip: Cover slides with aluminum foil to protect from light).
   a. 5 min
   b. 2x 15 min
   c. 5 min
2. Immerse slides in the DAPI solution for 5 minutes or add 100 µl of DAPI solution per slide.
3. Wash DAPI solution off slides 2x with PBS, for 5 minutes each.
4. Dry the edges of slides with a Kim wipe. Add Fluoromount™ to slides and apply coverslip.

*Adapted from Molecular Instruments HCR-FISH v3.0 system Protocol for FFPE Tissue Sections. https://www.molecularinstruments.com/
