## Supplementary File 3 for "Optimizing the hybridization chain reaction-fluorescence in situ hybridization (HCR-FISH) protocol for *Pleurodeles waltl*"

**Supplementary File 3**
**Step-by-Step Workflow Manual for Optional In-silico Workflow for Probe Design**

This manual describes four computational steps: NCBI transcript selection, BLASTX screening, Synteny validation, and Probe design. For clarity, these steps are organized into three main stages and using publicly available web interfaces, tools, and reference databases.

#### 3.1.1. Target Transcript Selection via Screening for conserved regions

*(Step 1)* Identify target transcript in NCBI.

1. Open [NCBI Genome Data Viewer]() page.
2. Search your gene of interest under "Search assembly".
3. Check how many transcripts the gene has by clicking the drop-down under "transcript". If there is more than one transcript for your gene, then proceed to the next step. If not, proceed to the synteny section.
4. Click the green double-sided arrow to reveal all the gene transcripts available for your gene of interest.

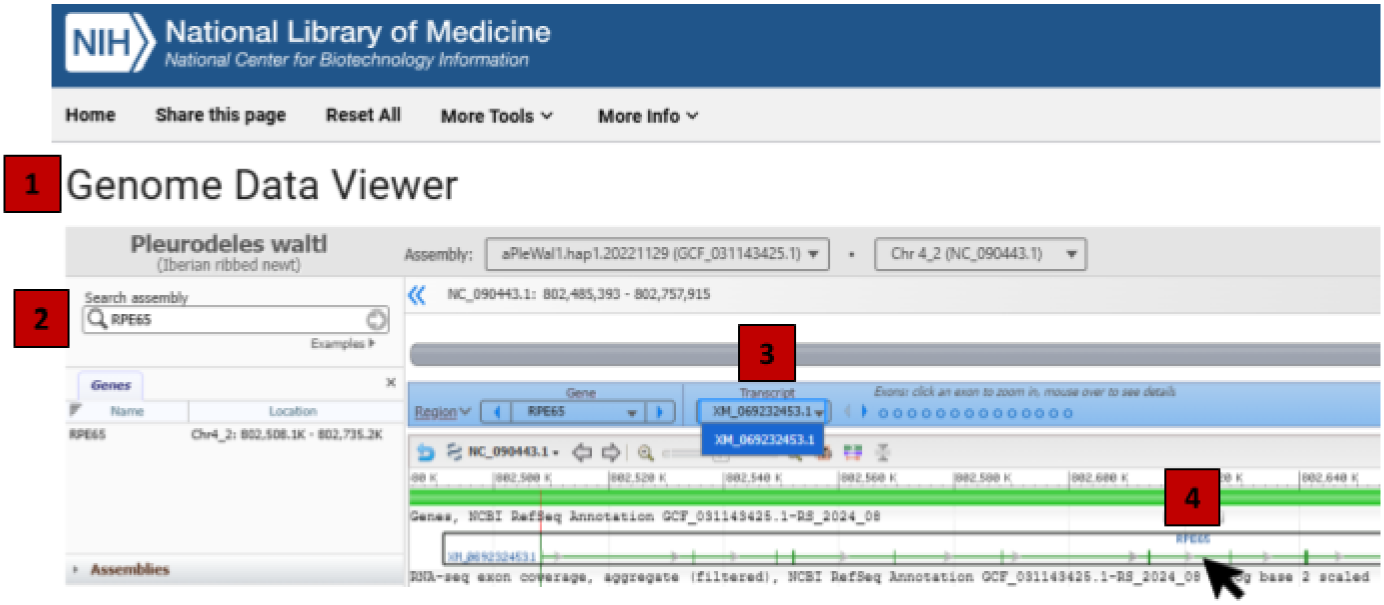


5. Pick the longest gene transcript with introns that you are interested in (purple) and hover your cursor over the transcript to see menu options to download the FASTA file.
   a. **Green** = gene
   b. **Purple** = spliced mRNA or complementary DNA (cDNA)
      i. If it starts with **NM**, these are curated NCBI reference transcripts.
      ii. If it starts with **XM,** these are predicted NCBI reference transcripts.
      iii. If you hover over the transcript, it will display its general information.
         - GeneID is under Links & Tools, which to open the NCBI Gene record for your gene of interest.

- From the table of contents, you can access the *NCBI Reference Sequences (RefSeq)*, which will contain a link to the associated UniProt gene information.
- On that page, the NCBI Orthologs section can be used for a rapid orthology check, but not at the isoform-specific level, because these assignments are gene-centered 1:1 orthology calls.

c. **Red** = spliced coding DNA sequence (CDS), which does not include the UTRs In this section, you may also see NP (known RefSeq protein record) and XP (model RefSeq records produced by prediction).

***Prioritize using full cDNA (mRNA sequence) because UTRs can differ between isoforms.**

*(Step 2)* Conserved-region screening using BLASTX.

6. Open [NCBI BLASTX]() page for translated nucleotide-to-protein searches.
7. Copy the name of the transcript ID or the whole sequence into the "enter accession number(s), gi(s), or FASTA sequence(s)" white box under the "enter query sequence" section.
8. Under "choose search set", click "add an organism" and add human and *Xenopus laevis* as organisms to blast against.
9. Click "Blast".

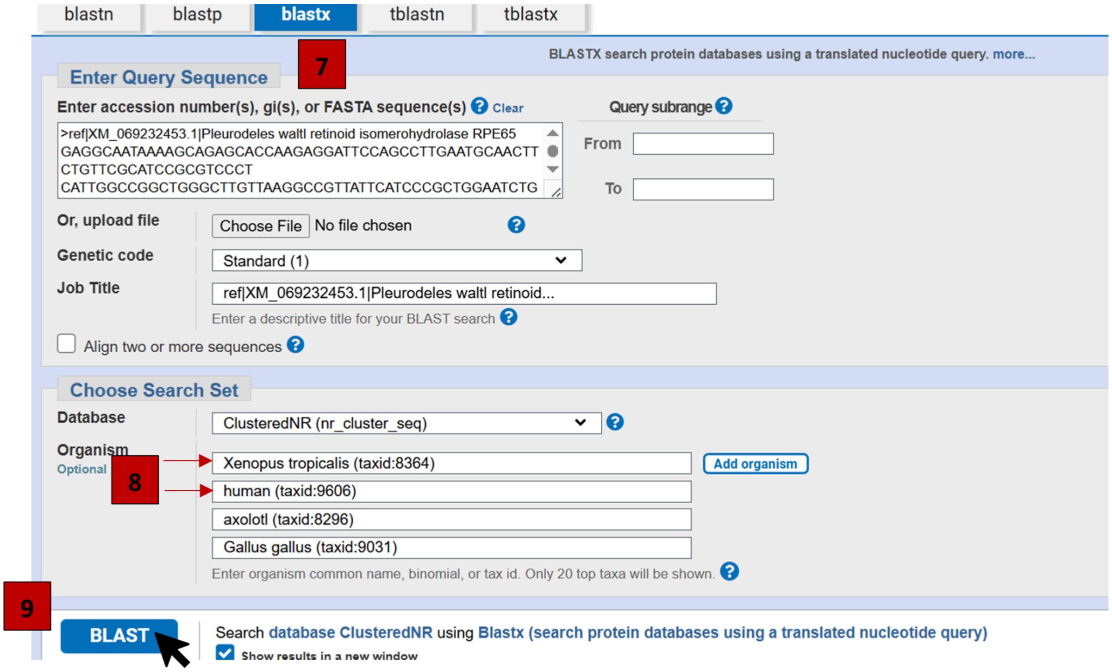


10. On "descriptions" result page, check scores for these three parameters:
    a. P value (E value) should be close to 0
    b. Percentage Identity should have higher values; it usually means the sequences are more similar (> 80%).
    c. Query Cover* with high numbers usually mean a stronger match of the target sequence with the aligned hit (>80%).

*Using mRNAs, the query cover can drop because of UTRs. So, a query cover below 50% with high percent identity may still be acceptable. For this reason, repeat the BLAST search using the CDS to obtain a more reliable estimate of query coverage (Appendix 1).

11. Under the “graphic summary” tab, verify if the query sequence covers the domains identified as important for the gene.
12. Repeat this with other transcripts if available to ensure that you are using the best transcript for probe design. Compare the 3 parameters listed above along with query sequence domain coverage.
13. Download the FASTA file corresponding to the selected transcript, or record the transcript ID (e.g., XM_069232453.1).

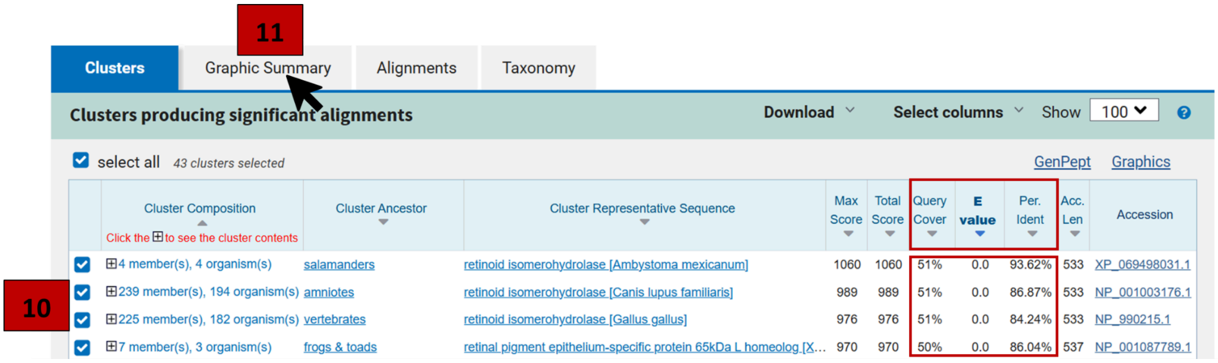


#### 3.1.2. <u>Orthology confirmation via synteny validation and paralog discrimination</u>

*(Step 3)* Confirm the expected ortholog by combining synteny visualization (MGV) with phylogenetic ortholog inference (SHOOT). Note: Checking for synteny (the conserved order of genes along a chromosome) helps verify that the target gene is truly orthologous to its counterpart in well-annotated species (e.g., frog, human). This ensures that the probe is being designed for the correct gene and not a paralog or a misassembled sequence.

1. Go back to the NCBI Genome Data Viewer for your gene and hover your cursor over the protein sequence (red double-sided arrow) for the transcript you selected.

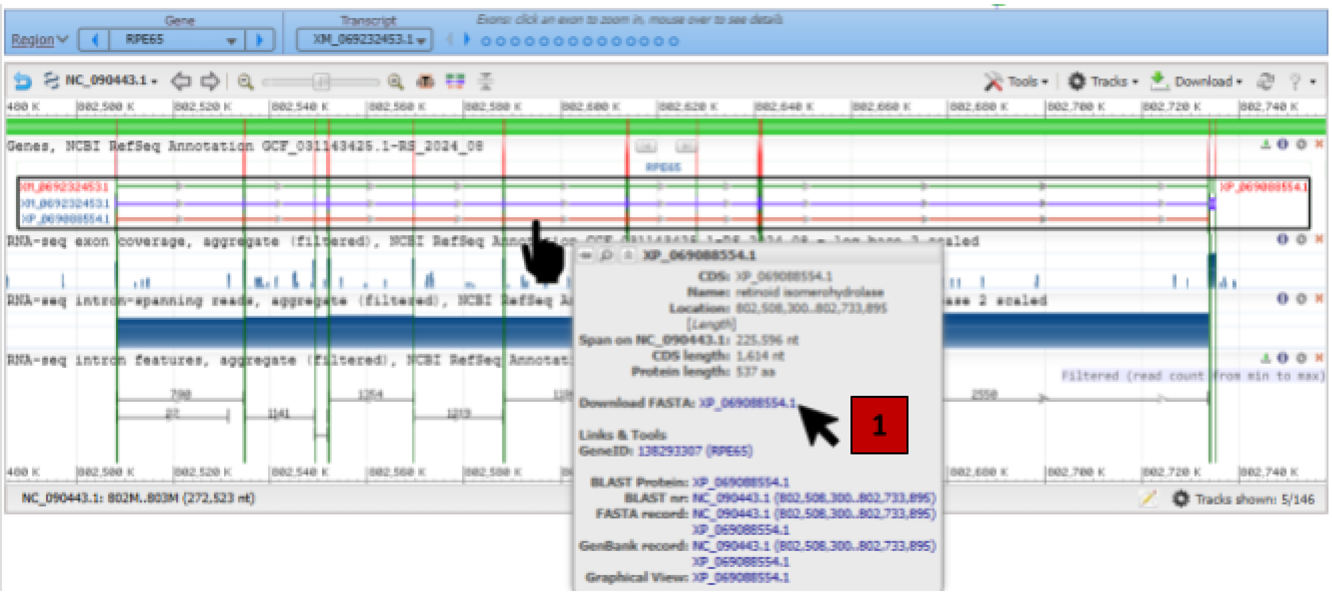

2. Download the FASTA File and copy the protein sequence, together with the gene name into [SHOOT](#) page. This tool compares your protein sequence to a database of gene families and provides you with a phylogenetic tree with your query sequence grafted into it. It helps us find the orthologs of our gene in other species.

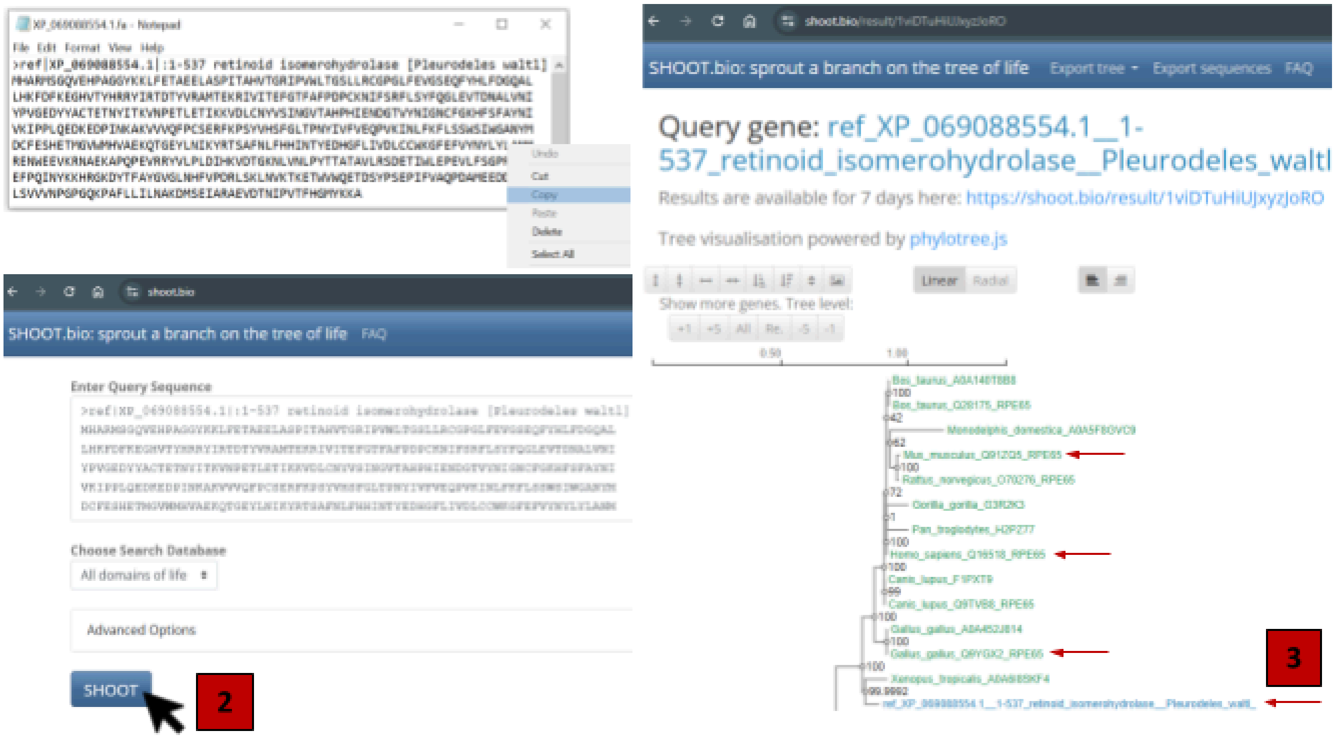


3. Check if your query sequence is placed within a phylogenetic tree to find sister branches (and related with the expected gene family), this can be taken as a support for orthology in candidates. In this manual, the **>95%** is used as a practical guide based on the SHOOT benchmark, rather than as a universal cutoff. For example, in our results, "*Xenopus_tropicalis_A0A6I8SKF4*" and "*ref_XP_069088554.1*" were among the strongest matches and are therefore reasonable candidate orthologs, with 99,9% bootstrap support. (If the tree looks inconsistent, try a different transcript, and look at synteny tools)
   *When several isoforms are available, the canonical or curated isoform should be used for comparison.
4. Use the [Multi Genome Viewer](#) (MGV) as another method of checking for synteny. This tool aligns several species genomes and makes it easier to check for the conservation of DNA sequences. You may also use [Xenbase](#) to check for synteny with Xenopus, specifically.
5. Compare the genes next to your gene of interest in MGV and compare to the genes next to your gene of interest in the NCBI Genome Data Viewer. If they have at least one gene next to each other, you can be more confident that we have the correct sequence for our gene of interest.
6. Once you have checked for synteny with our gene of interest, proceed to the next step.

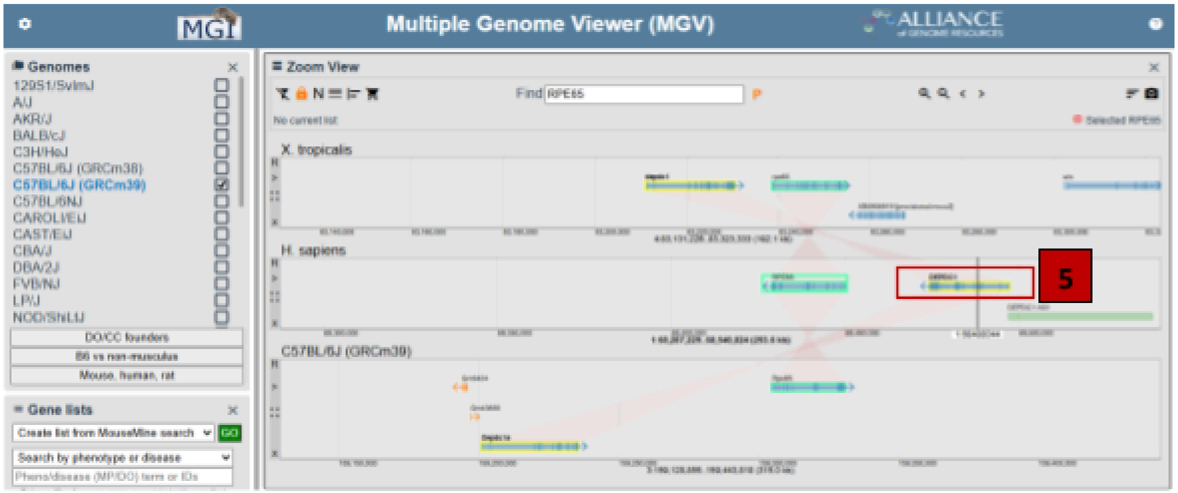


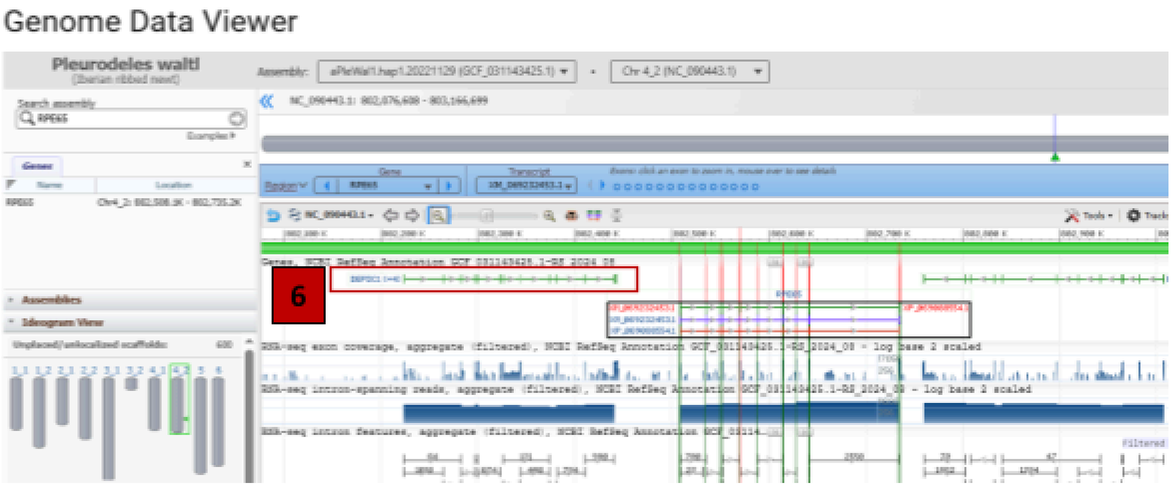


#### 3.1.3. <u>Probe design for IDT ordering</u>

*(Step 4)* Probe design tool generates split-initiator probe pairs targeting the transcript candidate, with an optional *BLASTn* filtering to reduce predicted off-target hybridization.

1. Access the probe design tool from this GitHub repository: [https://github.com/DelRioTsonisLab/HCR-probes-Tool.git](https://github.com/DelRioTsonisLab/HCR-probes-Tool.git)

   Review the *“Run in Google Colab (recommended)”* section to configure the notebook in your Google Drive. No additional code or package installation is required, as all dependencies are handled within the notebook. A detailed step-by-step guide is provided below and within the notebook for clarity.

2. In Google Colab, run the notebook sections in order (click the ▶️ “Run icon” and wait for each cell to finish ✅ before moving on): I. STORAGE SELECTION (2 cells), II. SETUP (6 cells), III. UPLOAD DATA (1 cells), and IV. START THE HCR PROBE MAKER (3 cells).

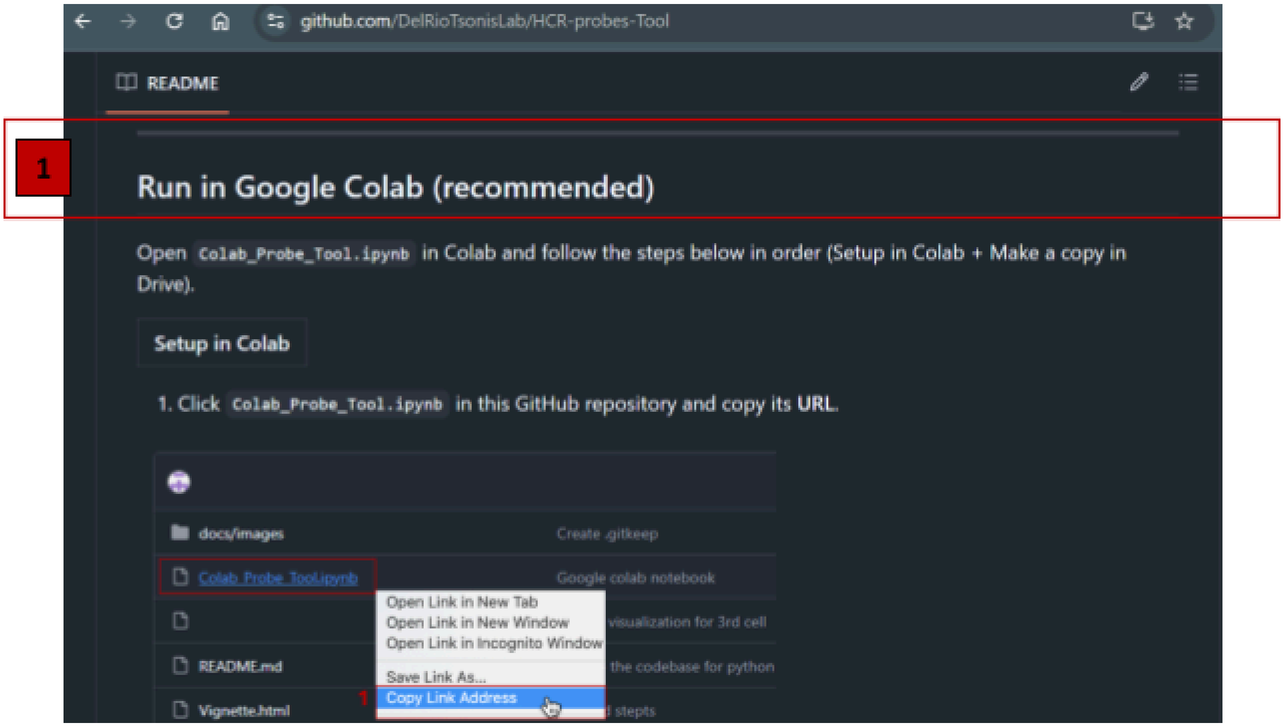


3. In STORAGE SELECTION section, click run to choose a storage option (the first cell ▶️), to know where FASTA file and outputs will be stored, if you select:

    a. 🔘*Google Drive (persistent)*, your FASTA file(s) and the exported .xlsx output can be saved persistently and reused in future sessions (you can later select Use existing files in the UPLOAD DATA section).

    b. 🔘*Local runtime /content (temporary)*, files are not preserved between sessions; you will typically need to select 🔵*Upload FASTA file(s) now* option in the UPLOAD DATA section each time you run the notebook.

   And, click run and wait until storage is selected (the second cell ▶️), to confirm that storage is selected.

4. In SETUP section, click run (the cell ▶️), to setup the working directory, download code resources, and install all dependencies.

   Run the SETUP block sequentially (cell-by-cell) or use the option to execute the entire SETUP section in one run ▶️ (if available in your interface).

5. In UPLOAD DATA section, click run to upload or locate FASTA file (the cell ▶️), then choose your data source from:
    a. 🔘*Use existing files* (if the FASTA is already present in the configured data folder), or
    b. 🔘*Upload FASTA file(s) now* (to add new FASTA files).

Select 🟦*Overwrite existing files with same name* option, only if you intend to replace a file that already exists with an updated version.

Then click ➢***Continue*** button to confirm that the notebook detects the cDNA FASTA file(s) of interest [**Note 1**] successfully.

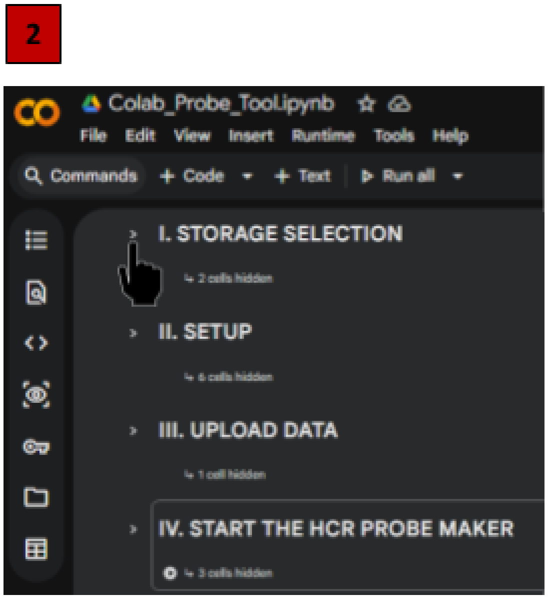


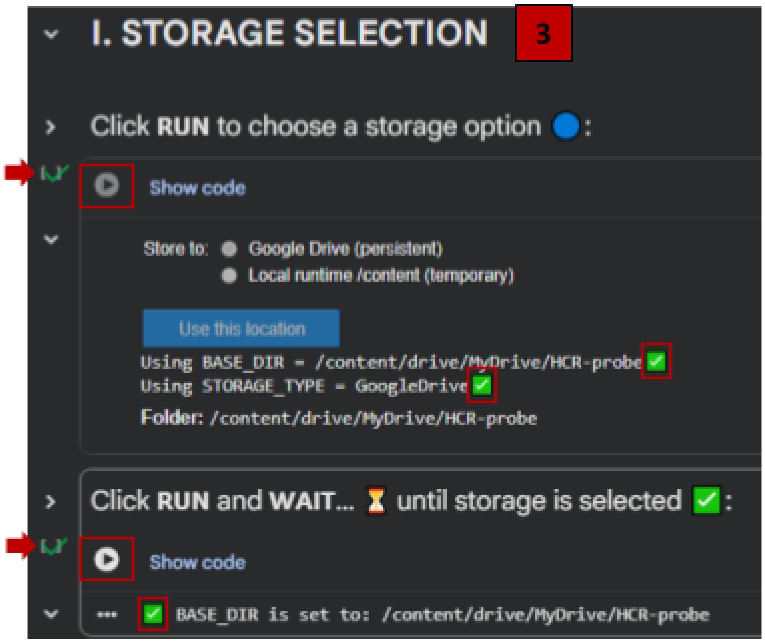


6. Under START THE HCR PROBE MAKER section, click run to initialize the environment (the first cell ▸ ). After it finishes, click run to set input parameters (the second cell ▶️), then it launches the interface.
7. Inside the ***Custom Probe Design Parameters*** interface, complete core information:
   a. Enter *Gene Symbol*: (e.g., Pax6)
   b. *Enter your full sense-strand cDNA (5'-3')* in one of two ways:

      i. Paste the target transcript sequence directly, or

      ii. Check ☑️*Select transcript from FASTA,* to load a sequence from a FASTA file.

      When ☑️*Select transcript from FASTA* is enabled, your FASTA file will appear under "*Select cDNA FASTA file*". If you need to choose a different file, click ➢***Change***. Otherwise, click ➢***Get Transcripts*** to list all transcripts available for the Gene Symbol entered above.

   c. To select a specific transcript, scroll to the 🖱️ *Transcript* dropdown, choose your transcript of interest, and click ➢***Use transcript*** to load it. Confirm that the chosen transcript is loaded and verify the reported length (number of bp) is correct before proceeding.
   d. *Choose the hairpin you will use to amplify*: (e.g. B3)

e. *How many bases from the 5’ end of the Sense RNA before starting to hybridize?* (e.g. 60)
f. *Change the tolerated homopolymer lengths for poly(A/T) and poly(C/G)*: (e.g. 5)

To increase BLAST specificity, check the settings under ***Default parameters***:

g. Perform BLASTn on*:* ☑️*Potential Probes*
h. Do you want to eliminate probes that appear in low quality BLAST outputs? ☑️*Drop low quality probes*
i. Do you want to display detailed BLAST outputs? ☑️*Show detailed blast outputs*
j. Do you want to display chosen parameters in output? ☑️*Display parameters chosen (The only option already checking)*

Click ➤***Store Inputs*** button to save the chosen settings.

8. Finally, click run to generate probes and export file (the third cell ▶️), it will prompt for user input:
    a. *Enter the number of probes that you want to test*: Type the total number of available probes reported in the output (e.g., “There are X possible probes for your sequence with the chosen parameters” then type X).
    b. *Enter the number of the probe group with which you wish to proceed*: Type the probe-set group number (e.g., 0) corresponding to the column of [start,stop] positions along the cDNA, then press ⏎ **Enter**.
9. After completion, an order-ready Excel file named <*Gene Symbol*>.xlsx is generated in the selected working directory. Open the file and review the **Pool name** and **Sequence** columns to verify the oligos and select the probe set to order.
   The same fields are also shown in the notebook output; scroll to the “Pool name, Sequence” section to inspect each oligo set.

10. Upload the resulting .xlsx file to the IDT ordering interface (or other **oligonucleotide synthesis** provider). Place the oligo order using the IDT Oligo Pool Entry page.
    After ordering, continue with the HCR-FISH procedure (see S1 and S2 files, experimental sections) and imaging to document probe signal and localization. Imaging steps are not detailed in this manuscript.

### Appendix 1. CDS-based validation

The coding sequence (CDS) is used as a secondary validation step to confirm conservation of the protein-coding region and support target identity. This is used as a confirmation step when the levels of alignment transcript are weak or ambiguous.

**5.1** Hover over the red CDS section of the transcript you want. Then click “Download FASTA” to get the CDS sequence in FASTA format. You can also click “BLAST Protein”. This will take you to the NCBI BLASTP page, where the accession number is filled in automatically, so you can continue with the next steps more easily.

Genome Data Viewer

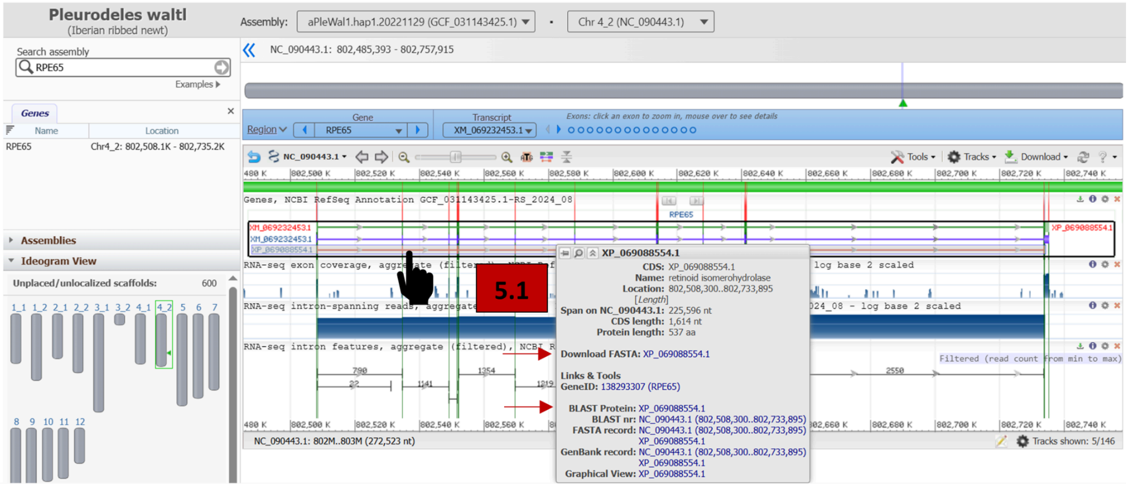


**6.1-9.1** You can either copy the accession number or paste the CDS FASTA sequence directly into the BLAST page. For a first search, you can keep the default database, such as ClusteredNR, and add the organism(s) you want to compare with your sequence. Then click “BLAST” to run the search.

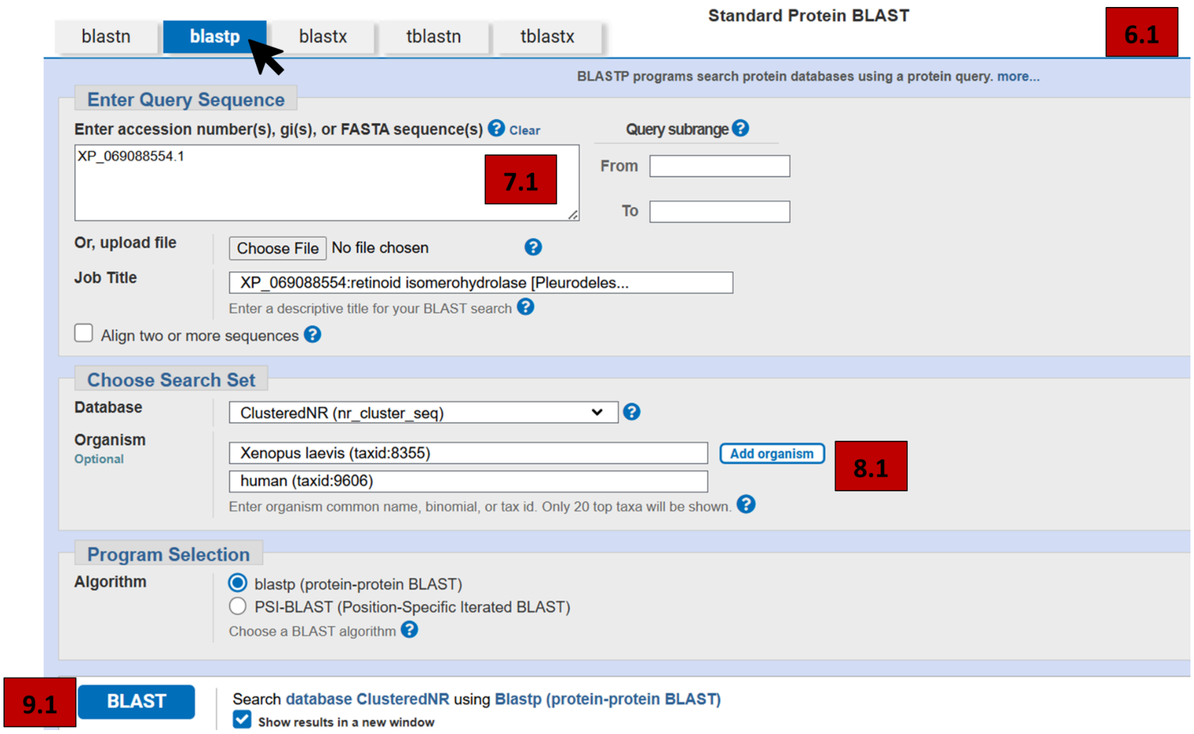

**10.1** On the "Clusters" results page, check for three parameters: P value (E value) should be close to 0, Percentage Identity should be >80%, Query Cover should be >80%. For a strong candidate, we started looking at those thresholds to exclude false positive sequences.

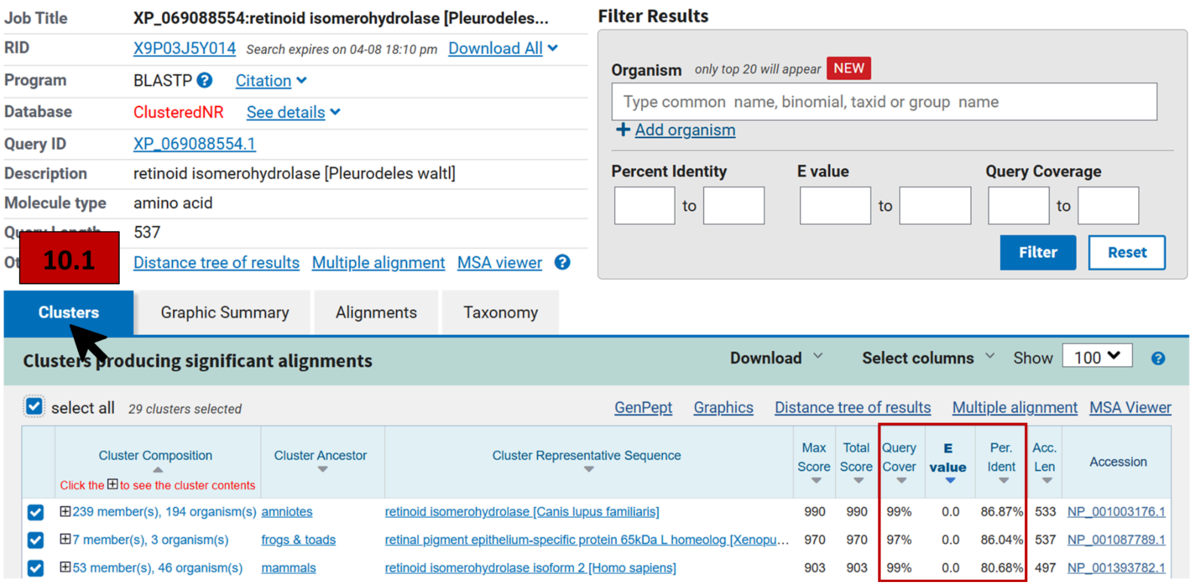


**11.1** In addition, under the "Graphic Summary" tab, check whether your query sequence covers the domain(s) that are important for your gene. You can use the "Specific hits" information to see where conserved domain(s) is located and compare it with your query sequence.

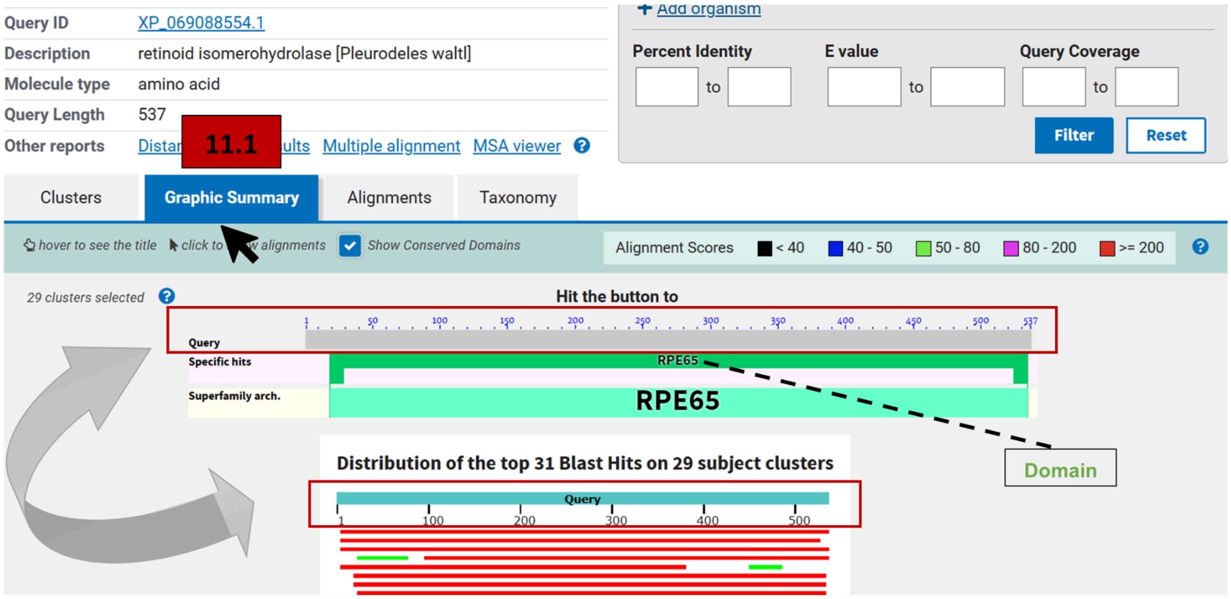


*In BLASTX results, the same query may appear with labels such as RF+2] or [RF-1], etc., which indicate different reading frames used to translate the nucleotide query during the search.

**12.1-13.1** You can repeat this process for each sequence you want to check and then download the CDS sequence (of the transcript you decided) to use in the next steps to validate your target sequence.
